# Ribosolve: Rapid determination of three-dimensional RNA-only structures

**DOI:** 10.1101/717801

**Authors:** Kalli Kappel, Kaiming Zhang, Zhaoming Su, Wipapat Kladwang, Shanshan Li, Grigore Pintilie, Ved V. Topkar, Ramya Rangan, Ivan N. Zheludev, Andrew M. Watkins, Joseph D. Yesselman, Wah Chiu, Rhiju Das

## Abstract

The discovery and design of biologically important RNA molecules is dramatically outpacing three-dimensional structural characterization. To address this challenge, we present Ribosolve, a hybrid method integrating moderate-resolution cryo-EM maps, chemical mapping, and Rosetta computational modeling, and demonstrate its application to thirteen previously unknown 119-to 338-nucleotide protein-free RNA-only structures: full-length *Tetrahymena* ribozyme, hc16 ligase with and without substrate, full-length *V. cholerae* and *F. nucleatum* glycine riboswitch aptamers with and without glycine, *Mycobacterium* SAM-IV riboswitch with and without S-adenosylmethionine, and computer-designed spinach-TTR-3, eterna3D-JR_1, and ATP-TTR-3 with and without AMP. Blind challenges, prospective compensatory mutagenesis, internal controls, and simulation benchmarks validate the Ribosolve models and establish that modeling convergence is quantitatively predictive of model accuracy. These results demonstrate that RNA-only 3D structure determination can be rapid and routine.

## Main Text

RNA molecules fold into intricate three-dimensional structures to perform essential biological and synthetic functions including regulating gene expression, sensing small molecules, and catalyzing reactions, often without the aid of proteins or other partners (*1, 2*). It is estimated that more than eighty percent of the human genome is transcribed to RNA, while just 1.5 percent codes for proteins, but our knowledge of RNA structure lags far behind our knowledge of protein structure (*3*). The Protein Data Bank, the repository for three-dimensional structures, currently contains fewer than 1,400 RNA structures, compared to ~143,000 protein structures. Accurate all-atom models of RNAs could dramatically enhance our understanding of functional similarities between distantly related RNA sequences, enable visualization of the conformational rearrangements that accompany substrate and ligand binding, and accelerate our ability to design and evolve synthetic structured RNA molecules. However, the conformational heterogeneity of RNA molecules, particularly in the absence of protein partners, challenges conventional structure determination techniques such as X-ray crystallography and NMR (*4, 5*). Even when such techniques are applied, the process is often laborious, time-consuming, and requires extensive construct-specific optimization and, typically, publications have reported only one or two 3D RNA structures at a time ((*4, 6*) and references therein).

Single-particle cryo-EM may provide an orthogonal approach to RNA structure determination. Recent advances in the technique have enabled high-resolution structure determination of proteins and large RNA-protein complexes that previously could not be solved with X-ray crystallography or NMR (*7, 8*). However, it has been widely assumed that most functional noncoding RNA molecules that are not part of large RNA-protein complexes are either too small or conformationally heterogeneous to characterize with cryo-EM. To date there is only one published sub-nanometer resolution cryo-EM map of an RNA molecule produced without protein partners, a 9 Å map of the 30 kDa HIV-1 dimerization initiation signal (DIS) (*9*). However, to reconstruct atomic coordinates into this map, detailed supplementary NMR measurements were required. In addition to its cost in time and specialized equipment, this approach may not be generally applicable due to the size limitations of NMR, motivating us to search for alternative approaches to complement cryo-EM. Here, we present Ribosolve, a pipeline integrating cryo-EM with biochemical RNA secondary structure determination and computational three-dimensional structure modeling (Fig. 1A).

**Fig. 1.**
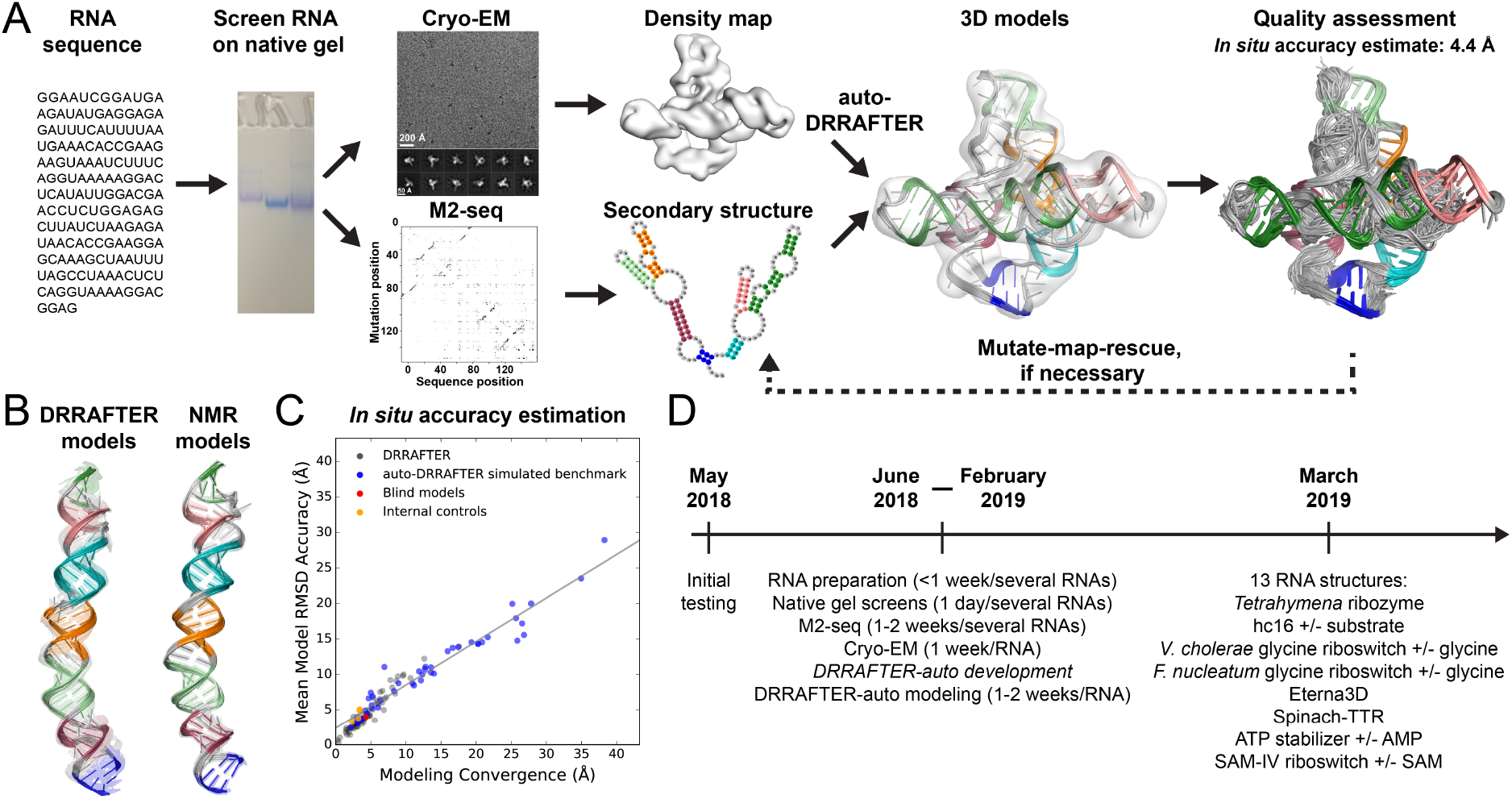
The Ribosolve pipeline. RNA structure determination through M2-seq, cryo-EM and computational modeling. (A) RNA samples are prepared and screened on a native gel to check for the formation of sharp bands. M2-seq experiments are performed to elucidate the RNA secondary structure. Cryo-EM elucidates the global architecture of the RNA. The M2-seq-based secondary structure and cryo-EM map are used to build all-atom models with auto-DRRAFTER. Model accuracy is predicted from the overall modeling convergence. Per-residue-convergence (top) and real-space correlation between the map and model (bottom) can help identify regions of uncertainty in the models. (B) Blind HIV-1 DIS Ribosolve models (left) and NMR models (right). (C) Mean model RMSD accuracy over the top ten scoring models versus modeling convergence for models after each round of auto-DRRAFTER for the benchmark on simulated maps (blue), DIS blind models (red), DRRAFTER models (gray) (*10*), and the automatically built *V. cholerae* and *F. nucleatum* glycine riboswitch Ribosolve models (internal controls, orange). Points representing models from the final round of auto-DRRAFTER modeling for each system are colored red. (D) Timeline for Ribosolve structure determination in this study.

### Computational modeling accurately builds RNA coordinates into cryo-EM maps

We hypothesized that combining moderate-resolution cryo-EM with computational three-dimensional structure modeling methods might enable rapid, simple, and general RNA structure determination. To rigorously evaluate this hypothesis, we performed a blind test using the 9 Å resolution DIS map. Prior to the publication of the HIV-1 DIS structure, K.K., A.M.W., and R.D. built all-atom models into the 9 Å map using a modified version of DRRAFTER, a recently developed Rosetta computational tool for modeling RNA coordinates into moderate-resolution density maps (see Methods) (*10*) (Fig. 1B). K.Z. and W.C. kept the coordinates derived from NMR restraints hidden while predictions were being made. In addition to building DRRAFTER models, we used the previously established linear relationship between model RMSD accuracy and modeling convergence, defined as the average pairwise RMSD across the top ten scoring DRRAFTER models, to predict that the models would have mean RMSD accuracy of 4.3 Å (gray points, Fig. 1C; and see below) (*10*). Indeed, the blind DRRAFTER models agreed well with the NMR models, with mean RMSD accuracy of 4.0 Å (Fig. 1B), suggesting that combining cryo-EM with computational modeling could provide a route to rapid and simple RNA structure determination.

### A benchmark set of RNA molecules with previously unknown structures

To more broadly benchmark this cryo-EM-DRRAFTER pipeline, we selected a set of eighteen functionally diverse RNA molecules ranging in size from 62 to 388 nucleotides (20 – 125 kDa) (Fig. 1D). The complete structures of these molecules were unknown (see below), and up to fifteen were expected to have well-defined 3D structures based on the functions they are known to perform. These RNAs broadly fall into three functional classes: ribozymes, riboswitches and computationally designed RNAs. The ribozymes include the L-21 ScaI ribozyme from *Tetrahymena thermophila*, which catalyzes a splicing reaction (*11*), the *in vitro* selected hc16 RNA ligase in an apo state and after ligation of an RNA substrate to its 5′ end (“hc16 product”) (*12*), and the 24-3 ribozyme, an *in vitro* selected RNA polymerase, which replicates short RNA sequences (*13*). The riboswitches, which regulate gene expression by sensing specific small molecule substrates, include the *V. cholerae* and *F. nucleatum* glycine riboswitch aptamers with and without glycine (*14*), a metagenomic SAM-IV riboswitch with and without S-adenosylmethionine (SAM) (*15*), and a metagenomic downstream peptide (glutamine-II) riboswitch with glutamine (*16*). The computationally designed synthetic RNA constructs include ATP-TTR-3, a stabilized aptamer for ATP and AMP (*17*), both with and without AMP; spinach-TTR-3, a stabilized version of the spinach aptamer (*17*), which fluoresces when bound to DFHBI (3,5-difluoro-4-hydroxybenzylidene imidazolinone) (*18*); and a molecule developed with a prototype 3D design interface in the Eterna online game, eterna3D-JR_1 (see Fig. S1) (*19*). Parts of the glycine riboswitches and *Tetrahymena* ribozyme were previously structurally characterized and therefore served as internal positive controls (*20-22*). However, all of these molecules contained substantial portions of unknown structure, in some cases after decades of attempts with prior methods (*21, 23*). As negative controls, we also included three molecules that were not expected to adopt well-defined three-dimensional structures: the human small Cajal body-specific RNA 6 (scaRNA6) and the human spliceosomal U1 snRNA, both of which function as part of larger RNA-protein complexes (*24, 25*) without any known RNA-RNA tertiary contacts and are therefore unlikely to be highly structured in the absence of their protein binding partners; and the human Retinoblastoma 1 (RB1) 5′ UTR, which has previously been shown to adopt multiple secondary structures (*26*).

### Cryo-EM easily resolves the global architectures of RNA molecules

After screening all RNA molecules on a native gel (Supplementary Results, Fig. S2, Fig. 1A), we used single-particle cryo-EM to try to resolve their three-dimensional architectures. As expected, we did not resolve the global architectures of the three negative controls, which were predicted not to form well-defined tertiary structures (Fig. 2P, Q, R). Additionally, the downstream peptide riboswitch, the smallest molecule in our benchmark set (62 nucleotides), and the 24-3 ribozyme, ran as a smear on a native gel, suggesting that they would exhibit substantial conformational flexibility in standard buffer conditions for *in vitro* RNA assembly (Fig. S2). Indeed, we did not resolve the global architectures of these molecules. To our surprise, we were able to resolve the global architectures of all remaining thirteen molecules in our benchmark set using standard cryo-EM experimental procedures with 1-2 days of image acquisition and standard data processing methods for each RNA molecule (Fig. 2, Methods). The final map resolutions ranged from 4.7 Å to 14 Å (Table S1, Fig. S3) and exhibited several distinctive characteristics of RNA molecules. All density maps contained rod-like shapes with dimensions concordant with RNA helices (~20 Å diameter; scale bars in Fig. 2). Major grooves are visible in ten of the maps (~12 Å wide, Fig. 2, blue arrows) and minor grooves are visible in seven of the maps (~17 Å wide, Fig. 2, red arrows). The relative sizes of the resolved molecules varied in accordance with the lengths of the RNA sequences; the *Tetrahymena* ribozyme map is the largest, while the SAM-IV riboswitch map is the smallest (Fig. 2S). Additionally, these maps exhibit several more complex features that were not present in the previously determined 9 Å HIV-1 DIS map (Fig. 2S) (*9*). The maps for each distinct RNA sequence reveal intricate 3D folds with tertiary features such as pockets and holes idiosyncratic to each RNA. To obtain more detailed insights into these rich features, we sought to model atomic coordinates into the density maps.

**Fig. 2.**
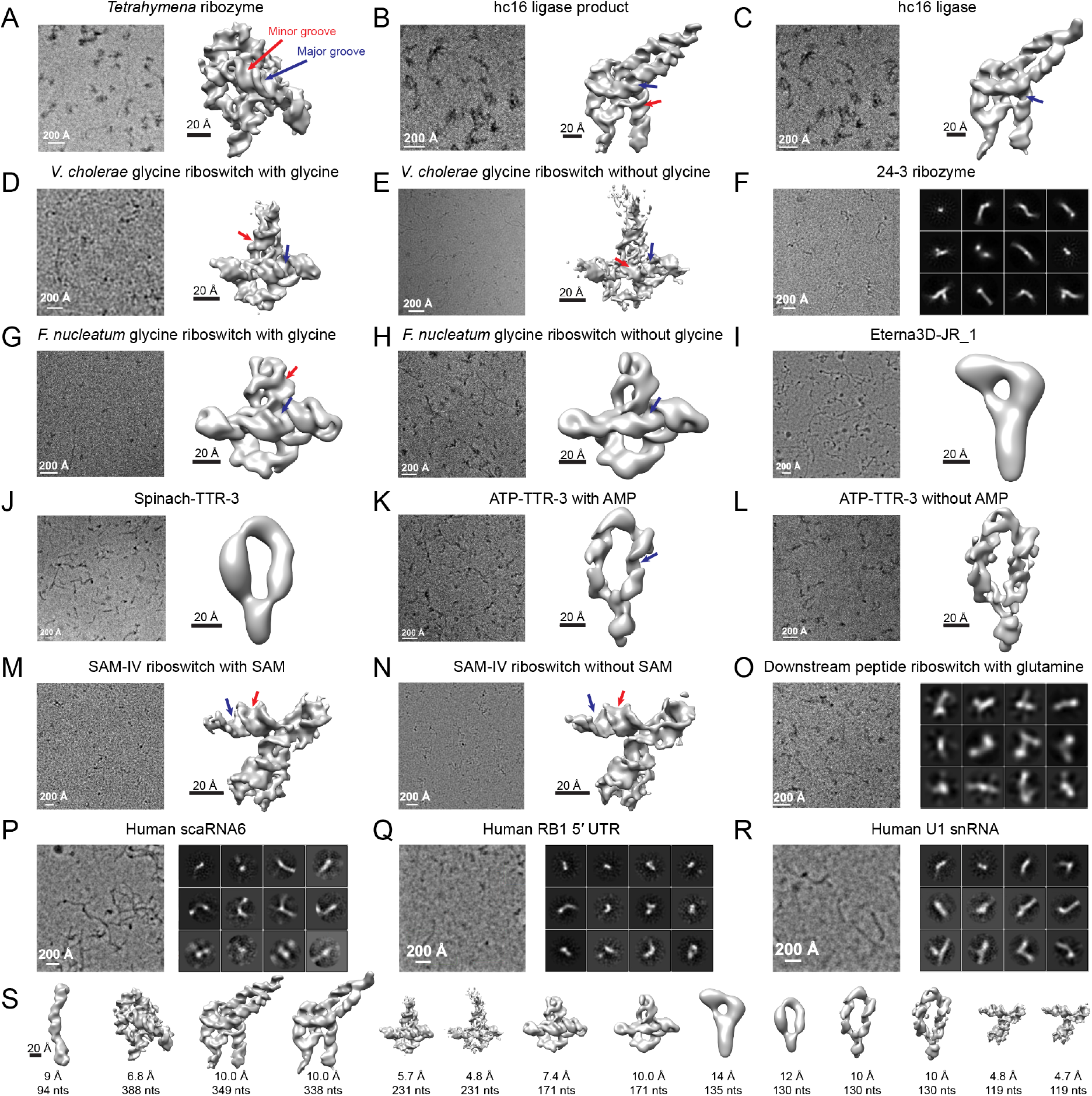
Cryo-EM resolves the global architectures of RNA molecules. Cryo-EM micrographs (left) and 2D class averages or 3D reconstructions (right) for (A-O) ribozymes, riboswitch aptamers, and synthetic RNA nanostructures, arranged in order of RNA size, largest to smallest. (P-R) Micrographs (left) and 2D class averages (right) for negative controls. (S) All cryo-EM maps of RNA molecules produced without protein partners to date. From left to right: HIV-1 DIS map (*9*) and all maps from this work shown in the same order as in (A-O). Map resolutions and the number of nucleotides in the RNA constructs are noted below each map. All maps are shown at the same scale.

### auto-DRRAFTER automatically models RNA coordinates into cryo-EM maps

Though our cryo-EM maps resolved the global architectures of the RNA molecules, they were not of sufficiently high resolution to enable manual atomic model building. Our blind tests on the HIV-1 DIS map suggested that we could instead apply computational modeling to build accurate models. However, DRRAFTER, the Rosetta method that we used for the DIS test case, was developed specifically for large RNA-protein complexes and requires initial manual setup, which may challenge and bias the modeling process particularly for smaller RNAs without protein partners (*10*). To address this problem, we developed auto-DRRAFTER to automatically model coordinates into moderate-resolution maps of RNA molecules (Fig. S4). Briefly, starting from an RNA sequence, secondary structure, and cryo-EM map, auto-DRRAFTER uses graph decomposition techniques (*27*) to automatically place at least one RNA helix into the density map and then samples conformations for the rest of the RNA through fragment-based RNA folding (Fig. S4C-J). To elucidate or confirm RNA secondary structures, we used mutate-and-map read out by next-generation sequencing (M2-seq) (*28*) (Supplementary Text; Fig. S5 – Fig. S7). The Rosetta low-resolution and full-atom RNA potentials used for scoring structures were augmented with a score term that rewards agreement with the density map (*29, 30*). Analogous to hybrid modeling methods in protein structure prediction (*31, 32*), iterative modeling was performed in several rounds. Hundreds to thousands of models are built in each round, then automatically checked region-by-region for structural consensus across top scoring models. Regions with sufficient consensus were then kept fixed in the next round. This automated process is continued until the entire structure can be confidently built. Final refinement was carried out in two independent cryo-EM maps generated from separate halves of the cryo-EM data. Additional details are provided in the Methods. Tests using simulated density maps of eight RNAs of known structure suggest that auto-DRRAFTER models are accurate (Fig. S8, Fig. S9, Table S2, Supplementary Text). Additionally, these tests demonstrate that auto-DRRAFTER modeling convergence, defined as the average pairwise RMSD over the top ten scoring models, is correlated with model RMSD accuracy (r^2^ = 0.95), suggesting that convergence can be used to predict model accuracy (blue points, Fig. 1C.).

### Thirteen all-atom models from cryo-EM, M2-seq, and auto-DRRAFTER

The Ribosolve pipeline enabled us to build all-atom models for all thirteen RNAs for which we had acquired cryo-EM maps with coordinate RMSD accuracies ranging from 3.3 Å to 6.3 Å as estimated by modeling convergence (Fig. 3, Table 1, Table S3, Fig. 1C). As further tests of model accuracy, we performed additional consistency checks for each of the thirteen Ribosolve structures (Supplementary Text), including comparison of automatically built models to crystallographic conformations of sub-structures confirming RMSD accuracies of 4.9 Å and 3.3 Å for *F. nucleatum* and *V. cholerae* glycine riboswitches (Fig. S10), mutate-map-rescue experiments to confirm novel secondary structure rearrangements for hc16 (Fig. S11-Fig. S13), and comparisons of structures to functional and biochemical data for hc16, the SAM-IV riboswitch, ATP-TTR-3, spinach-TTR-3, and Eterna3D-JR_1 (Fig. S10, Fig. S14). The coordinate accuracy length scale of 3.3-6.3 Å is finer than the typical distance between backbone atoms in consecutive nucleotides and, historically, has been sufficient to attain non-trivial insights into whether RNAs of similar function have similar folds, how RNA structures respond to binding partners, and whether designed RNA structures fold as predicted (*23, 25, 33, 34*). Each of the thirteen Ribosolve models revealed at least one of these kinds of insights. These results are described briefly below.

**Table 1.** Estimated accuracy of Ribosolve models^a^

| System | No. nts | Final no. particles used for 3D reconstruction | Map resolution (Å) | Estimated accuracy<sup>a</sup> (Å) |
| --- | --- | --- | --- | --- |
| *Tetrahymena* ribozyme | 388 | 74,621 | 6.8 | 3.9, 3.9, 3.9 |
| hc16 product | 349 | 29,191 | 10.0 | 6.3, 6.2, 6.4 |
| hc16 | 338 | 21,263 | 10.0 | 6.2, 6.2, 6.1 |
| *V. cholerae* glycine riboswitch with glycine | 231 | 193,317 | 5.7 | 3.9, 3.9, 3.9 |
| *V. cholerae* glycine riboswitch apo | 231 | 230,891 | 4.8 | 3.3, 3.3, 3.2 |
| *F. nucleatum* glycine riboswitch with glycine | 171 | 35,578 | 7.4 | 3.4, 3.3, 3.4 |
| *F. nucleatum* glycine riboswitch apo | 171 | 20,269 | 10.0 | 3.5, 3.6, 3.4 |
| Eterna3D-JR_1 | 135 | 51,690 | 14.0 | 3.9, 3.9, 3.9 |
| Spinach-TTR-3 | 130 | 67,474 | 12.0 | 4.2, 4.2, 4.2 |
| ATP-TTR-3 with AMP | 130 | 39,136 | 10.0 | 3.7, 3.6, 3.9 |
| ATP-TTR-3 apo | 130 | 71,045 | 10.0 | 3.7, 3.7, 3.6 |
| SAM-IV riboswitch with SAM | 119 | 225,303 | 4.8 | 3.6, 3.6, 3.6 |
| SAM-IV riboswitch apo | 119 | 260,244 | 4.7 | 3.7, 3.7, 3.6 |

<sup>a</sup>Accuracy is estimated from the convergence, defined as average pairwise RMSD of the top ten scoring models built into half map 1 vs. models built into half map 2 (first number), models built into half map 1 (second number), and convergence of models built into half map 2 (third number).

**Fig. 3.**
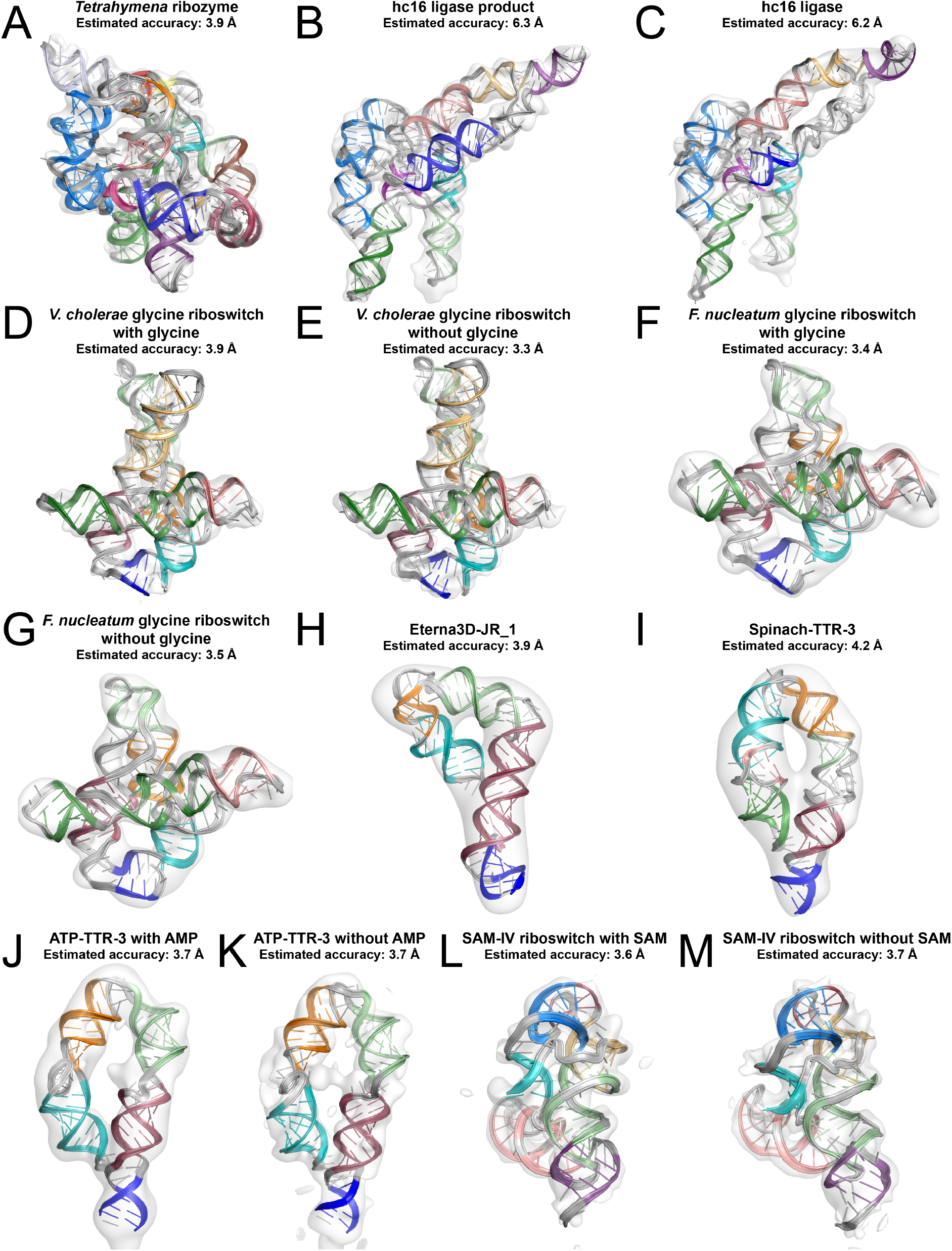
RNA structures determined by the Ribosolve pipeline. (A-M) Top scoring auto-DRRAFTER models in the cryo-EM maps and estimated accuracies based on modeling convergence.

### Glycine riboswitches from different species adopt nearly identical folds

The *F. nucleatum* and *V. cholerae* glycine riboswitches each contain two glycine aptamers that interact through tertiary contacts to form a butterfly-like fold (Fig. 3D-G, Fig. 4A). For both RNAs, we see evidence of two features predicted from previous computational analysis, a P0 stem (Fig. 3D-G, blue) and kink-turn formed between the 5′ ends of each molecule and the linker between each molecule’s two glycine aptamers (*35*). In addition to the clear structural homology between the two glycine riboswitches, the two aptamers within each riboswitch also adopt nearly identical folds (Fig. 4A-B). Furthermore for both riboswitches, the apo (ligand-free) states closely resemble the holo (ligand-bound) states (Fig. 3D-G). This invariance in tertiary fold has been observed previously for other natural riboswitches, but was missed for these glycine riboswitches because sequences previously used for structure determination were over-truncated (*20, 22*).

**Fig. 4.**
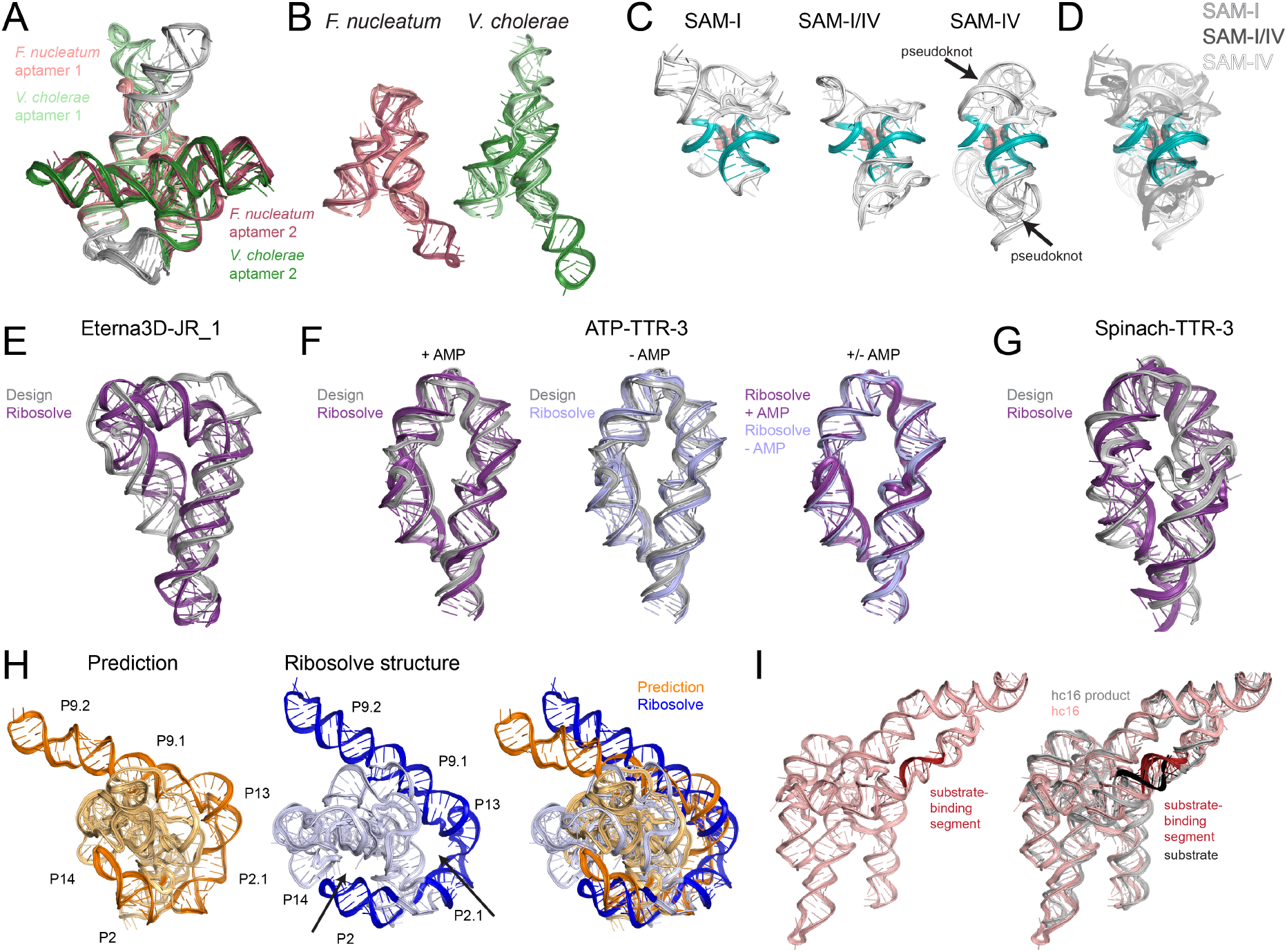
Functional insights from Ribosolve models. (A) Overlay of *V. cholerae* and *F. nucleatum* glycine riboswitches with glycine. (B) Overlay of both glycine aptamers from the *F. nucleatum* and *V. cholerae* structures. (C-D) Structural homology between the SAM-IV riboswitch and SAM-I and SAM-I/IV riboswitches. (C) The SAM-I crystal structure (*37*), the SAM-I/IV crystal structure (*36*), and the SAM-IV Ribosolve model, and (D) an overlay of all three structures, with peripheral elements shown as gray transparent cartoons. SAM is shown as transparent red spheres. (E) The computationally designed eterna3D-JR_1 structure (gray) overlaid with the Ribosolve model (purple). (E-G) Comparisons of Ribosolve structures and the computationally designed models for (E) Eterna3D-JR_1, (F) ATP-TTR-3 with and without AMP, and (G) Spinach-TTR-3. (H) The overall architecture of the previously predicted model of the *Tetrahymena* ribozyme (*38*) qualitatively matches the Ribosolve structure, with peripheral elements wrapping around the core. The Ribosolve structure contains holes (arrows in middle panel) not present in the predicted structure. (I) Comparison of the hc16 Ribosolve structures without substrate (pink) and with ligated substrate (gray). The substrate-binding segment and substrate are highlighted in red and black, respectively.

### Distinct peripheral architectures support a conserved core structure across multiple classes of SAM riboswitches

Ribosolve reveals that the SAM-IV riboswitch adopts a complex fold with two pseudoknots (Fig. 3L, M). Comparison of this structure with previously solved crystal structures of SAM-I and SAM-I/IV riboswitches reveals that the tertiary structure is substantially rewired across the three classes of SAM riboswitches (Fig. 4C), but the core structures are nearly identical in all three molecules (Fig. 4D) (*36, 37*). These observations are consistent with previously hypothesized homology based on secondary structure (*36*). Like the glycine riboswitches, the apo and holo SAM-IV riboswitch structures are highly similar (Fig. 3L, M).

### Assessing the accuracy of computationally designed RNA molecules

Beyond its application to determining structures of natural RNAs, Ribosolve structures enabled rapid validation or falsification of four synthetic RNA structures. First, the Eterna3D-JR_1 Ribosolve structure provides valuable feedback about the accuracy of this design. Eterna3D-JR_1 was computationally designed to adopt a triangular conformation, however, the computational design does not closely match the Ribosolve structure (RMSD = 15.7 Å; Fig. S15, Fig. 4E). The ATP-TTR-3 and spinach-TTR-3 embed the AMP aptamer and spinach RNA, respectively, into clothespin-like scaffolds, seeking to pre-organize the aptamer structures to enhance their ligand binding affinities (*17*), analogous to natural riboswitch aptamers including the glycine and SAM riboswitches solved above (Fig. 3D-G, L-M). Indeed, our structures of ATP-TTR-3 with and without AMP and spinach-TTR-3 confirm this pre-folding into the computer-designed tertiary structures within error (RMSDs of 4.2 Å, 4.3 Å, and 7.7 Å, respectively) (Fig. 4F-G, Fig. S15). Additionally, the ATP-TTR-3 Ribosolve structures with and without AMP are very similar (mean RMSD = 2.7 Å), further supporting the hypothesis that minimal conformational rearrangement occurs upon ligand binding (Fig. 4F). We envision that the Ribosolve pipeline could be used to rapidly evaluate many designed synthetic structures to accelerate the design-build-test cycle of RNA nanotechnology and medicine.

### The complete global architecture of the *Tetrahymena* ribozyme and rearrangements of core elements in the hc16 ligase

Ribosolve structures of our two largest molecules offered rich comparisons to literature predictions and to each other. Through the Ribosolve pipeline, we resolved the first complete experimental 3D structure of the *Tetrahymena* ribozyme, a paradigmatic RNA enzyme discovered 35 years ago and manually modeled 20 years ago (*38*) (Fig. 3A). Peripheral elements of the ribozyme, P2, P2.1, P9.2, P9.1, P13, P14, are resolved for the first time here; these elements wrap around the core of the ribozyme, to complete a ring around its catalytic core, qualitatively substantiating a decades-old prediction (Fig. 4H), while also revealing large holes in the structure that were not previously predicted (Fig. 4H, arrows) (*38*). We expected the architecture of the hc16 ligase to be very similar to the *Tetrahymena* ribozyme because hc16 was evolved *in vitro* from a random library that contained the P4-P6/P3-P8 domain of the *Tetrahymena* ribozyme as a constant scaffold region (*12*). Instead, the hc16 Ribosolve structure exhibits a unique extended conformation that has not yet been observed in other ribozyme structures (Fig. 3B, C). In both the structures without substrate and with ligated substrate, the active site is at the center of the molecule and the substrate-binding segment is positioned like the string of a bow (Fig. 4I). This rearrangement of the hc16 ligase from its ‘parent’ *Tetrahymena* ribozyme, while unexpected, provides a rationalization for paradoxical observations in prior sequence conservation data for hc16, is consistent with new chemical mapping data for the RNA (Supplementary Text; Fig. S11-Fig. S13), and provides a particularly powerful example of how unbiased structure determination enabled by Ribosolve can falsify and refine structure-function-homology relationships.

## Discussion

The Ribosolve pipeline combines recent advances in cryo-EM, M2-seq biochemical analysis, and Rosetta auto-DRRAFTER computer modeling to accelerate three-dimensional RNA structure determination. Using this method, we solved the structures of thirteen RNA molecules, including riboswitches, ribozymes, and synthetic RNA nanostructures, over the timescale of months (Fig. 1D). A gauntlet of tests, including a blind challenge, internal controls, and simulation benchmarks, validate and confirm that Ribosolve models have coordinate accuracies of 3-6 Å. For every one of the thirteen Ribosolve models determined here, the resolution is sufficient to reveal global tertiary structural features, to detect structural rearrangements upon target binding, to confirm or falsify hypotheses of homology between RNA classes, or to efficiently validate or falsify structure predictions. Use of the Rosetta auto-DRRAFTER tool, rather than manual coordinate modeling, not only accelerates model building into maps but also enables reliable prediction of model accuracy. Important frontiers for Ribosolve include inference of RNA conformational ensembles rather than single dominant conformations (*39*); automatic refinement of RNA secondary structures during 3D model building, which may obviate M2-seq experiments; and systematic tests of RNAs smaller than 100 nucleotides with varying numbers of tertiary contacts. Nevertheless, Ribosolve, even in its current form, provides at least an order-of-magnitude acceleration in our ability to experimentally infer 3D RNA structures at nucleotide resolution. Our study suggests that it should now be possible to determine structures of the thousands of noncoding RNA domains that have been proposed to adopt well-defined tertiary folds without partners (*40*).

## Acknowledgments

We thank members of the Das lab for useful discussions; members of the Rosetta community for discussions and code sharing; M. Summers for permission to perform the HIV-1 DIS blind modeling challenge; V. Kosaraju, and J. Nicol for helping to develop the Eterna3D interface; JR for designing the eterna3D-JR_1 construct through the Eterna3D interface; and the Warsaw Do Science Club for feedback on this work. Calculations were performed on the Stanford Sherlock cluster.

## Author contributions

K.K., R.D., and W.C. conceptualized and designed the research. K.K. prepared the RNA samples for cryo-EM. K.Z., Z.S., S.L., and G.P. collected and analyzed the cryo-EM data. K.K. and V.V.T. collected and analyzed the M2-seq data. W.K. collected the mutate-map-rescue data. W.K. and K.K. analyzed the mutate-map-rescue data. K.K. developed, implemented, and tested the computational approach with input from R.R. and R.D. K.K., A.M.W., and R.D. performed the blind DIS modeling. K.K. performed modeling for all other RNA systems. I.N.Z. prepared the 24-3 ribozyme RNA for cryo-EM and performed functional validation. K.K. and R.D. wrote the manuscript with input from all authors.

## Data and materials availability

Cryo-EM maps will be deposited in the EMDB. Models will be deposited in the PDB. M2-seq and mutate-map-rescue data will be deposited in the RMDB. The auto-DRRAFTER software will be freely available to academic users as part of the Rosetta software package.

