## Supplemental Materials for "Ribosolve: Rapid determination of three-dimensional RNA-only structures"

**This PDF file includes:**

Materials and Methods
Supplementary Results
Figs. S1 to S21
Tables S1 to S5
Caption for Table S6

**Other Supplementary Materials for this manuscript include the following:**

Table S6

### Materials and Methods

RNA preparation

DNA templates for all molecules except *Tetrahymena* ribozyme were prepared by PCR assembly of DNA oligonucleotides designed with Primerize (*1*) and purchased from Integrated DNA Technologies (Table S6). PCR reactions contained 1× HF buffer (New England Biolabs (NEB)), 0.2 mM dNTPs, 4 μM end primers, 40 nM middle primers, and 40 U/mL Phusion Polymerase. Reactions were first denatured at 98 °C for 30 s. Then for 35 cycles, they were denatured at 98°C for 10 s, annealed at 67 °C for the *V. cholerae* glycine riboswitch and 65°C for all other RNAs, and incubated at 72 °C for polymerase extension for 30 s. Finally, reactions were incubated at 72 °C for 10 min. DNA templates were purified with AMPure XP beads (Beckman Coulter) following the manufacturer's instructions. RNA transcription reactions for all RNAs except *Tetrahymena* ribozyme contained 0.2 μM DNA template, 40 mM Tris·HCl, pH 8.1, 25 mM $MgCl_2$, 3.5 mM spermidine, 0.01% TritonX-100, 40 mM DTT, 4% PEG 8000, 3 mM NTPs, and 5 U/μLT7 RNA polymerase (NEB). Reactions were incubated at 37 °C for 4 h. RNAs were purified with Zymo RNA Clean and Concentrator columns (Zymo Research) following the manufacturer's instructions.

The DNA template for the *Tetrahymena* ribozyme was PCR amplified from the pT7L-21 plasmid (*2*) using the primers listed in Table S6. The reaction contained 1× HF buffer (New England Biolabs (NEB)), 0.2 mM dNTPs, 4 μM primers, 40 pg/μL plasmid, and 40 U/mL Phusion Polymerase. The reaction was performed as described above except with an annealing temperature of 69°C and an extension time of 60 s. The DNA template was purified as described above. The transcription reaction contained 0.2 μM DNA template, 40 mM Tris·HCl, pH 8.1, 25 mM $MgCl_2$, 3.5 mM spermidine, 0.01% TritonX-100, 40 mM DTT, 4% PEG 8000, 3 mM NTPs, and 7.5 U/μLT7 RNA polymerase (NEB). The reaction was incubated at 37 °C for 1 hour. The RNA was then purified by ethanol precipitation as follows. 1/10 × volume of 3 M Na-acetate, pH 5.2, and 1.0 μL 15 mg/mL GlycoBlue (Thermo Fisher Scientific) was added to the transcription reaction and mixed. 3.3 × volume of chilled 100% ethanol was then added, then reactions were placed on dry ice for 20 minutes. The reactions were then spun down in a tabletop centrifuge at maximum speed for 30 minutes, then the supernatant was removed and discarded. The pellet was then washed twice by adding 500 μL 70% chilled ethanol, centrifuging for 10 minutes at maximum speed, and finally removing the supernatant. The pellet was then dried for 5 minutes and resuspended in RNase-free water. Approximately 10 μL of loading buffer containing 10 mM EDTA, 0.1% xylene cyanol, 0.1% bromophenol blue in 95% formamide was then added to the reaction. The *Tetrahymena* ribozyme RNA was then PAGE-purified, as follows. A 8% 29:1 acrylamid:bis, 7 M urea polyacrylamide gel was poured and allowed to set overnight. RNA containing loading buffer described above was loaded onto the gel and run at 25 W for 2 h. The RNA was visualized in the gel by brief exposure to a 254-nm UV lamp, held far from the gel to minimize RNA damage (*3*). RNA was eluted from the gel in RNase-free water overnight at 4 °C. RNA was then purified with Zymo RNA Clean and Concentrator columns (Zymo Research).

After preparing the RNA, we then confirmed that the three ribozymes in our benchmark set (*Tetrahymena* ribozyme, hc16, and 24-3) were catalytically active (Fig. S16, details below).

Native gels

All native gels were run using a BioRad Criterion Cell gel cassette. Polyacrylamide gels were cast by combining 15 mL of 8% or 12% 29:1 acrylamide:bis, in 10 mM $MgCl_2$, 67 mM

HEPES, 33 mM Tris, pH 7.2 solution with 150 μL 10% ammonium persulfate and 30 μL TEMED. After the gel polymerized, chilled buffer containing 33 mM Tris, 10 mM $MgCl_2$, 67 mM HEPES, pH 7.2 was added and the gel apparatus was placed in an ice bath in a 4 °C cold room. Folded RNA was prepared as follows. 1 μg of RNA was diluted to a volume of 6.4 μL, then incubated at 90°C for 3 minutes, then room temperature for 10 minutes. 0.8 μL of 500 mM Na-HEPES pH 8 and 0.8 μL of 100 mM MgCl2 was added, then the reaction was incubated at 50°C for 20 minutes, and finally for 10 minutes at room temperature. All RNAs were screened in the apo state. 2 μL of loading buffer containing 250 mM Na-HEPES pH 7.5, 5 mM EDTA pH 8, 50% glycerol, 0.05% xylene cyanol, and 0.05% bromophenol blue was then added and samples were then loaded into the gel immediately. The gel was run at 10 W for 1.5 hours. The temperature of the gel was monitored closely to avoid overheating. To visualize the RNA, the gel was submerged in Stains-All working solution (0.015% dye in 45% formamide) and placed on an orbital shaker for 25 minutes. The Stains-All solution was then removed and the gel was de-stained in water for 15 minutes. The water was then removed and the gel was imaged.

*Tetrahymena* ribozyme activity assay

To check that the *Tetrahymena* ribozyme was active, we first combined 150 nM RNA and 50 mM Na-HEPES, pH 8 and heated at 90°C for 3 minutes to denature the RNA. The solution was then cooled at room temperature for 10 minutes. 10 mM $MgCl_2$ was added and the solution was then incubated at 50°C for 30 minutes for folded RNA and at room temperature for 15 minutes for misfolded RNA. The solution was then cooled at room temperature for 10 minutes, then 500 μM GTP was added and the solution was incubated at room temperature for 5 minutes. 750 nM fluorescently labeled substrate (Cy5-CCCUCUAAAAA, purchased from IDT) was then added and the reaction was incubated at room temperature in the dark for 1 hour. All concentrations listed above are final. 2 μL of the reaction was then added to 10 μL of stop solution containing 20 mM EDTA and 90% formamide, to quench the reaction. The reaction was then heated at 90°C for 7 minutes. Immediately, 2 μL of the reaction was then loaded onto a 20% 29:1 acrylamide:bis, 7 M urea, 1 x TBE polyacrylamide gel, which had been pre-run at 10 W for 1 hour. The gel was run for 35 minutes at 15 W, and then imaged immediately with a Bio-Rad VersaDoc MP 4000 using the Cy5 detection setting.

hc16 activity assay

To check that the hc16 ribozyme was active, we performed ligation reactions as previously described (*4*), except that we used a fluorescently labeled substrate (Cy5-AUCUUACUU). Briefly, the hc16 RNA was incubated at 85 °C for 2 minutes and then 50 °C for 2 minutes, then diluted in buffer containing $MgCl_2$, KCl, spermidine, Na-HEPES, pH 7.5 and then incubated at 50°C for 10 minutes. The fluorescently labeled substrate was added and incubated in the dark at 50°C for 75 minutes. The final concentrations in the reaction were: 0.1 μM RNA, 0.2 μM substrate, 50 mM $MgCl_2$, 200 mM KCl, 5 mM spermidine, and 50 mM Na-HEPES, pH 7.5. To stop the reaction, 10 μL of 120 mM EDTA, 8 M urea solution was added to 10 μL of the reaction. The stopped reaction was then loaded onto a 20% 29:1 acrylamide:bis, 7 M urea, 1 x TBE polyacrylamide gel, which had been pre-run at 10 W for 1 hour. The gel was then run for 35 minutes at 15 W, then imaged immediately as described above.

24-3 activity assay

The RNA template for the 24-3 primer extension assay was prepared through PCR assembly and *in vitro* transcription as described above. A fluorescently labeled RNA primer was purchased from IDT. Primer and template sequences are listed in Table S6. The primer extension assay was performed as described previously (*5*). Briefly, reactions were conducted with final concentrations of 0.5 μM fluorescently labeled primer RNA, 0.5 μM template RNA, 0.5 μM 24-3 ribozyme, 0.05% Tween 20, 50 mM Tris pH 8.1, 200 mM $MgCl_2$, and 4 mM UTP, GTP, ATP, and CTP. Briefly, the RNA (in RNase free water) was first heated for two minutes at 80°C, then cooled at 17°C for ten minutes. Primer extension was initiated by adding chilled extension buffer (Tween 20, Tris, $MgCl_2$, NTPs) to the final concentrations listed above. The sample was mixed and allowed to extend at 17°C. The reaction was quenched by adding EDTA to a final concentration of 125 mM. Samples were then ethanol precipitated as described above, and resuspended in 70% formamide, diluted with TBE buffer. The samples were then run on a 20% 29:1 acrylamide:bis, 7 M urea, 1 x TBE polyacrylamide gel, then imaged for FAM fluorescence.

M2-seq experiments

M2-seq experiments were performed as previously described (*6*). Briefly, DNA templates for RNAs containing 5′ and 3′ buffer sequences were prepared by PCR assembly of DNA oligonucleotides designed with Primerize (*1*) and purchased from IDT (Table S6), except for the *Tetrahymena* ribozyme, which was PCR amplified from the pT7L-21 plasmid using the primers listed in Table S6. PCR products were checked by agarose gel electrophoresis. Reactions that produced only the desired product were purified with AMPure XP beads. Other reactions were purified by gel extraction using the Qiagen gel extraction kit following the manufacturer's instructions. Mutations were introduced in the templates through error-prone PCR as described previously (*6*). Again, reactions were purified with AMPure XP beads or by gel extraction. RNA transcription, RNA purification, DMS treatment, ethanol precipitation, reverse transcription, cDNA purification, and library preparation for sequencing were performed as described previously (*6*). Primers used for reverse transcription are listed in Table S6. M2-seq data analysis was performed with the pipeline available at https://github.com/ribokit/M2seq.

Briefly, multiplexed FASTQ sequence files were demultiplexed into sample-specific files using NovoBarcode (Novocraft Technologies). Demultiplexed, paired-end files were run through the ShapeMapper analysis pipeline (*7*). The ShapeMapper-derived mutation strings were converted to simple binary mutation files with `muts_to_simple.py`. These `.simple` files were then converted to `.rdat` 2D mutational profile files with `simple_to_rdat.py`. RDAT files were then analyzed by scripts in the Biers package (https://ribokit.github.io/Biers/) as follows. Z-scores incorporating DMS treated and untreated RNA for each sample were generated with `output_Zscore_from_rdat.m`. Secondary structure prediction was performed with ShapeKnots version 6.1 with the M2-seq data using the rna_structure MATLAB wrapper in the Biers software. 100 bootstrapping iterations were performed. The secondary structure predicted by ShapeKnots using all of the M2-seq data, but with base pairs with bootstrap probabilities less than 65% removed, was used for auto-DRRAFTER modeling (*8*).

Cryo-EM sample preparation, data collection, and image processing

RNAs were prepared as follows. RNAs were combined with Na-HEPES, pH 8 and incubated at 90°C for 3 minutes, then cooled at room temperature for 10 minutes. $MgCl_2$ and any ligands were added and the solution was incubated at 50 °C for 20 minutes (30 minutes for *Tetrahymena* ribozyme), then cooled at room temperature for 10 minutes. The final concentrations were 10 mM $MgCl_2$, 50 mM Na-HEPES, pH 8. The final RNA and ligand concentration were: 15 µM *Tetrahymena* ribozyme RNA; 15 µM hc16 product RNA; 15 µM hc16 RNA; 25 µM scaRNA6 RNA; 25 µM *V. cholerae* glycine riboswitch RNA and 100 mM glycine; 15 µM *V. cholerae* glycine riboswitch RNA; 21 µM RB1 5′ UTR RNA; 25 µM U1 snRNA RNA; 25 µM 24-3 RNA; 13 µM *F. nucleatum* glycine riboswitch RNA and 100 mM glycine; 17 µM *F. nucleatum* glycine riboswitch RNA; 20 µM Eterna3D-JR_1 RNA; 26 µM spinach-TTR-3 RNA; 27 µM ATP-TTR-3 RNA and 1 mM AMP; 30 µM ATP-TTR-3 RNA; 40 µM SAM-IV riboswitch RNA and 1 mM SAM; 45 µM SAM-IV riboswitch RNA; 40 µM downstream peptide riboswitch RNA and 10 mM glutamine.

Three microliters of the samples were applied onto glow-discharged 200-mesh R2/1 or R3.5/1 Quantifoil grids. The grids were blotted for 2-4 s and flash frozen in liquid ethane using a Vitrobot Mark IV (Thermo Fisher Scientific) with the chamber cooled to 4 °C and 100% humidity. The samples were screened using a Talos Arctica cryo-electron microscope (Thermo Fisher Scientific) operated at 200 kV. The *Tetrahymena* ribozyme, hc16 and hc16 product, RB1 5’ UTR RNA, U1 snRNA RNA, *F. nucleatum* glycine riboswitch with and without 100 mM glycine, Eterna3D-JR_1 RNA, spinach-TTR-3 RNA, ATP-TTR-3 with and without 1 mM AMP, downstream peptide riboswitch with 10 mM glutamine were imaged on a Talos Arctica cryo-electron microscope. *V. cholerae* glycine riboswitch RNA with and without glycine, SAM-IV riboswitch with and without SAM, and 24-3 RNA were imaged on a Titan Krios cryo-electron microscope (Thermo Fisher Scientific) with GIF energy filter (Gatan) Micrographs were recorded by EPU software (Thermo Fisher Scientific) with a Gatan K2 Summit direct electron detector, where each image was composed of 30 individual frames with an exposure time of 6 s.

All micrographs were motion corrected using MotionCor2 (*9*) and the contrast transfer function (CTF) was corrected using CTFFIND4 (*10*). All particles were autopicked using the NeuralNet option in EMAN2 (*11*), and further checked manually. Then, particle coordinates were imported to Relion (*12*), where the 2D classification was performed. Several rounds of 2D classification were performed to remove poor 2D classes. Initial 3D maps were generated in cryoSPARC (*13*) for RB1 5′ UTR RNA, U1 snRNA RNA, Eterna3D-JR_1 RNA, spinach-TTR-3 RNA, *V. cholerae* glycine riboswitch RNA with and without glycine, SAM-IV riboswitch with and without SAM, 24-3 RNA, and in EMAN2 for *Tetrahymena* ribozyme, hc16 with and without ligation product, *F. nucleatum* glycine riboswitch with and without glycine, ATP-TTR-3 with and without AMP, and downstream peptide with glutamine. 3D classification and final 3D refinement were performed in cryoSPARC for SAM-IV riboswitch with and without SAM and in Relion for *V. cholerae* glycine riboswitch RNA with and without glycine, Eterna3D-JR_1 RNA, spinach-TTR-3 RNA, *Tetrahymena* ribozyme, hc16 with and without ligation product, *F. nucleatum* glycine riboswitch with and without glycine, ATP-TTR-3 with and without AMP, and downstream peptide riboswitch with glutamine. An example of the data processing workflow is shown for the apo state of the *V. cholerae* glycine riboswitch in Fig. S17. Additional information about the data collection and image processing can be found in Table S1.

Auto-DRRAFTER pipeline

The inputs for the auto-DRRAFTER pipeline are an RNA sequence, secondary structure, and cryo-EM map. First, ideal A-form helices are built for all base paired regions of the structure. Helices are then placed in the density map as follows. The density map is first low-pass filtered to 20 Å, to identify “end nodes” or regions of the map that correspond to hairpin loops or helices formed between the very 5' and 3' ends of an RNA (Fig. S4B). Points are then placed in the density map using e2segment3d in EMAN2 using the command (*11*):

```
e2segment3d.py DENS_MAP --pdbout=ouput.pdb --
process=segment.distance:maxsegsep=18:minsegsep=15:thr=MAP_THR
```

where MAP_THR is provided by the user. A graph is then constructed from these points, where each point becomes a node, and nodes are connected by an edge if they are within 20 Å. For points that were initially connected to just one other node, additional edges are added to other points that are within 27 Å and have a "connecting density score" of greater than half of the average connecting density score for sets of points with connecting density scores greater than 0.02. The connecting density score is defined as the average of the density values at 5 equally spaced points between two points. "End nodes" are then defined as nodes that are connected to only one other point or nodes that are connected to multiple points for which the angle formed between a first neighboring point, the node of interest, and a second neighboring point is less than 0.87 radians, for all possible sets of two neighboring points (Fig. S4B). End nodes are also defined in the secondary structure by converting the secondary structure to a graph, where the helices, loops, and junctions are represented as nodes with edges between elements containing adjacent nucleotides, then identifying nodes that are connected to only one other node (Fig. S4A).

End nodes in the secondary structure are then mapped to end nodes in the density map (Fig. S4C). If more than two end nodes in the map, or more than 8 consecutive nucleotides that are not base paired in the input secondary structure, then just a single end node in the map is randomly selected for placement of an end node from the secondary structure. If there are exactly two end nodes identified in the map, then all possible mappings between secondary structure end nodes and density map end nodes are considered. For secondary structure end nodes that are hairpins, the 3D structure of the hairpin is modeled onto the adjacent helix through RNA fragment assembly in Rosetta (*14*) then the helix is fit into the density map at the location of the end node. This fitting is carried out by aligning a six base pair probe helix to the end node in the map and one of its neighbors (referred to as the neighboring map node below). The position of the helix is then optimized by randomly perturbing the probe helix through rigid body translations and rotations, keeping the distance between the C1' atom of the sixth base pair of the probe helix less than 15 Å or the starting distance between these points and keeping the distance between the C1′ atom of the sixth base pair of the probe helix and the neighboring map node greater than or equal to the initial distance between these points minus 8 Å. Each conformation is scored using the Rosetta `elec_dens_fast` score term (*15*). Conformations are accepted if the score of the current conformation is less than the score of the previous conformation. 5000 perturbations are attempted. The entire process is repeated 10 times and the best scoring placement is taken as the final probe helix placement. All helices from the secondary structure are then aligned to the optimized probe helix to generate the initial conformations for the auto-DRRAFTER simulations (Fig. S4C).

This process may result in several possible placements of helices within the density map. For each placement, the rest of the RNA is built with through RNA fragment assembly in Rosetta, keeping the placed helix fixed throughout the run (*14*). The low-resolution and all-atom Rosetta score functions are augmented with the `elec_dens_fast` score term to monitor agreement with the density map (*16*).

Modeling was then performed in several rounds (Fig. S4C-J). For each round, 1000 models (for the simulated benchmark) or 2000 models (for the best-case and automated models built into experimental maps) were built for each placement. The average pairwise RMSD (convergence) was then computed for the top ten scoring models across all helix placements. If the convergence was higher than 10 Å, then another round of modeling was performed. To set up the next round of modeling, for each helix placement, the convergence and average and minimum density scores for each node in the RNA, defined by the RNA secondary structure as described above, were computed over the top ten scoring models. Here, the average density score is defined as the average of the density values at the positions of each of the atoms within an element. The minimum density score is defined as the minimum of the density values at the positions of each of the atoms within an element. For each node, the average and minimum density scores were calculated in each of the top ten scoring models. The highest average density score was then compared to a threshold defined as 40% of the maximum density value in the density map. The minimum density score for the model with the highest average density score was also compared to this threshold. The convergence value was also compared to a threshold defined by the number of nucleotides in the node and whether it is a helix. The threshold was 7.0 Å for helices containing 3 or fewer base pairs, 10.0 Å for 4 base pair helices, 20.0 Å for 5 base pair helices, and 25.0 Å for helices containing 6 or more base pairs, and 4.0 Å for elements other than helices. If the convergence for the node was below the appropriate threshold and the average and minimum density scores were above the appropriate threshold, then the 3D conformation of this node was extracted from the model in which it had the highest density score and kept fixed for the subsequent rounds. Conformations were not taken from outlier models, defined as models for which $|\bar{C} - C_i| > 3\sigma$, where $C_i$ is defined for a model $i$ as the convergence when that model $i$ is removed from the top ten scoring models (i.e. convergence computed over the other nine models), $\bar{C}$ is the average of the $C_i$ values for the top ten scoring models and $\sigma$ is the standard deviation. Nodes that clash with extracted structures for any other node are removed.

When the convergence of the top ten scoring models across all helix placements dropped below 10 Å, the process described above was applied to the overall top ten scoring models (rather than the top ten scoring models for each helix placement) to identify regions of the structure that are well-converged and fit well in the density map, with the following differences (Fig. S4G-H). All conformations were taken from the top scoring model rather than the model with the best average density score. Additionally, the conformations taken from the models from the previous round were used as initial placements for the next round of modeling, but were allowed to move throughout the run. All other parameters remained the same in the subsequent round of modeling.

If the convergence does not drop below 10 Å and the average change in the convergence between the last three consecutive rounds is less than 1.5 Å after seven rounds, then subsequent rounds of modeling are not performed. Typically this is an indication that one of the initial assumptions of the modeling, such as the RNA secondary structure or initial helix placements, was incorrect and these inputs should be carefully reexamined.

For the final round of modeling, regions of the models that were well-converged and fit well in the density were again identified using the same procedure as for the previous round of modeling (Fig. S4I-J). These conformations were kept fixed during the fragment assembly stage of the modeling, but allowed to move along with the rest of the structure during the all-atom refinement stage. This last modeling round is performed in parallel in separate half maps, if available. The total numbers of modeling rounds performed for all RNAs described here are listed in Table 1, Table S2, Table S4. One round of modeling takes approximately one day on 50 × (number of helix placements) cores (Intel Xeon E5-2640v4 processors).

Auto-DRRAFTER benchmark on simulated maps

Starting with the database of nonredundant RNA PDB structures (release 3.39) solved to 4.0 Å resolution or better (*17*), we found all RNA structures of length between 100 and 450, removed structures with protein or DNA residues, and kept structures with single chains and limited missing residues. This yielded 25 structures, from which we selected a set of eight functionally and structurally diverse RNAs: THF riboswitch (PBD ID: 3SUX) (*18*), c-di-AMP riboswitch (PDB ID: 4QK8) (*19*), bacterial SRP Alu domain (PDB: 4WFL) (*20*), FMN riboswitch (PDB ID: 3F2Q) (*21*), SAM-I riboswitch (PDB ID: 4KQY) (*22*), *Tetrahymena* ribozyme P4-P6 domain (PDB ID: 1GID) (*23*), lysine riboswitch (PDB ID: 3DIL) (*24*), and the lariat capping ribozyme (PDB ID: 4P8Z) (*25*). Density maps were simulated at 10 Å resolution with EMAN2 using the following command:

```
e2pdb2mrc.py crystal_structure.pdb simulated_map.mrc --res=10.0
--center
```

We then built models with auto-DRRAFTER starting from the RNA sequences, secondary structures derived from the crystal structures, and simulated density maps. Fragments from homologous RNA structures were excluded during the fragment assembly stages of the auto-DRRAFTER runs with the following flags:

```
-fragment_homology_rmsd 1.2 -exclusion_match_type MATCH_YR
-exclude_fragment_files crystal_structure.pdb
```

where crystal_structure.pdb is the previously solved crystal structure. RMSDs between auto-DRRAFTER models and the crystal structure were calculated over all heavy atoms after alignment over all heavy atoms.

HIV-1 DIS blind modeling challenge

Models were built in five different ways using an early version of auto-DRRAFTER that required manually placing at least one helix in the cryo-EM density map and in which all modeling was performed in a single round (*26*). For the first set of models, the center helix was manually fit into the cryo-EM density map (chain A residues 257-262 and chain B residues 257-262), and the rest of the RNA was built through the fragment assembly procedure described here. We then built a second set of models starting from the best scoring model from the first set of models. During this run, all helices were allowed to move independently as rigid bodies and nucleotides in all junctions were allowed to move. During the final refinement stage, all nucleotides were allowed to move. For the third set of models, we started auto-DRRAFTER runs from chain A residues 244-276 and chain B residues 244-276 from the previously solved NMR model of HIV-1 DIS (PDB ID: 2D1A) (*27*), after first mutating it to the correct sequence. Throughout the run, helices were allowed to move as rigid bodies, though they were not allowed to dock independently (`-dock_each_chunk_per_chain false`), and nucleotides in junctions were also allowed to move. All nucleotides could move during the final refinement stage. For the fourth set of models, an initial conformation for chain A residues 248-270 and chain B residues 248-270 was taken from the first set of models. The remaining RNA was built with constraints for possible interactions that could occur within the other four RNA junctions. The constraints were based on configurations observed in initial stepwise Monte Carlo runs of these junctions without the density map (*28*). These constraints were also used to build the final set of models. These models were built using stepwise Monte Carlo with the density map.

All five sets of models were refined with a modified ERRASER protocol (*29*). Briefly, the entire structure was first minimized with restraints on the phosphate positions using the default ERRASER kinematics and also using the kinematics from the auto-DRRAFTER runs. The full structure was subsequently minimized again without restraints, then individual nucleotides were rebuilt using the standard ERRASER protocol, followed by a final round of minimization over the entire structure.

Auto-DRRAFTER models for experimental density maps

Automated models were built for all experimental density maps using the auto-DRRAFTER method as described above using the secondary structure derived from our M2-seq experiments. Additionally, we built a set of best-case auto-DRRAFTER models for these same systems. For this set of models, the M2-seq-based secondary structures were modified based on sequence covariation information and previously solved crystal structures. Additionally, specific elements of the RNA structures were initially placed in the density maps rather than using the automatic helix placement strategy described above. These elements were allowed to move from their initial positions during all rounds of modeling. For the ATP stabilizer with and without AMP and the spinach-TTR-3, the P1 helices were initially fit manually in the density map. Additionally, a model of the tetraloop-tetraloop receptor taken from a crystal structure of the P4-P6 domain of the *Tetrahymena* ribozyme (PDB ID: 1GID) was included as a rigid body (*23*). The conformation of the tetraloop-tetraloop receptor was refined during the final round of modeling. For the *F. nucleatum* glycine riboswitch with and without glycine, the initial conformation for residues 1-20, 25-71, and 80-158 was taken from a previously solved crystal structure (PDB ID: 3P49) (*30*), which was initially fit into the density map. During the runs, helices were allowed to dock independently as rigid bodies, and nucleotides in junctions were allowed to remodel. For the *Tetrahymena* ribozyme, the initial conformations for nucleotides 98-168, 174-209, 211-234, 240-258, 260-268, 270-276, 305-321, 324-325, 327-331, 405-406 were taken from a previously solved crystal structure (PBD ID: 1X8W) (*31*), which was initially fit in the density map. For the SAM-IV riboswitch with and without SAM, P5 (residues 87-91 and 99-102) was manually placed in the density map. Additionally, the structure of the SAM binding pocket (nucleotides 3, 7, 42, 63, 64, and 78) was taken from a previously solved crystal structure of the SAM-I riboswitch (PDB ID: 2GIS) (*32*). For models built into the map solved without SAM, these nucleotides were allowed to remodel after the initial round of modeling. For models built into the maps solved with and without SAM, these nucleotides were allowed to move during the final all-atom refinement during the final round of modeling. For the *V. cholerae* glycine riboswitch with and without glycine, the initial conformation of residues 141-155, 162-187, and 193-224 was taken from a previously solved crystal structure (PDB ID: 3OWI) which was initially fit into the density map (*33*). The conformations of these nucleotides were refined during the final round of modeling. For hc16 and the hc16 product, we built a preliminary model by first fitting P6a, P6b, P5b, the tetraloop-tetraloop receptor (residues 95-100, 167-173, and 192-197), and the part of P1 formed between the substrate and the ribozyme into the density map for the first conformation of the hc16 product. The rest of the RNA was then built with auto-

DRRAFTER while keeping these regions fixed. The helices and the tetraloop-tetraloop receptor (residues 95-100, 167-173, and 192-197) from the best scoring preliminary model were used as an initial conformation for auto-DRRAFTER runs for hc16 and the first conformation of the hc16 product. These elements were allowed to move as rigid bodies throughout the runs and were allowed to move during the final round of refinement. For the second conformation of the hc16 product, the complete best scoring preliminary model was used as an initial conformation for auto-DRRAFTER runs. Regions that were not in helices were allowed to remodel. We built models for hc16 and both conformations of the hc16 product with four possible secondary structures in the regions of uncertainty (dashed lines, Fig. S7): P10-ext and P11, P10-ext and alt-P11, P12 and P11, P12 and alt-P11. Models with alt-P11 and either P10-ext or P12 were built as described above. Only a final round of modeling was performed for models with P11, starting from the second to final round of models built with alt-P11 and either P10-ext or P12, but with the region around P11/alt-P11 completely rebuilt. The final set of models for these systems contains the top scoring models for each of these possible secondary structures. For eterna3D-JR_1, we did not have any additional information beyond the M2-seq-based secondary structure and cryo-EM map, so we did not build a best-case set of models. Ligands were not included in the automated or best-case modeling. The final round of modeling (described above), including refinement into half maps, was performed for all best-case models.

Auto-DRRAFTER error estimates and model validation

To estimate model error, we calculated the auto-DRRAFTER modeling convergence, defined as the average pairwise RMSD over the top ten scoring auto-DRRAFTER models. Models were not aligned before computing RMSDs. For models built into half maps, the convergence was defined as the average pairwise RMSD between the top ten scoring models refined into each of the half maps. For models built into a single map, convergence can be calculated in Rosetta with the following command:

```
drrafter_error_estimation -s {models} -rmsd_nosuper
```

For models built into separate half maps, convergence can be calculated with the command:

```
drrafter_error_estimation -sgroup1 {models1} -sgroup2 {models2}
-rmsd_nosuper
```

where models1 and models2 are the top ten scoring models refined into the two separate half maps.

Modeling convergence correlates with model accuracy ($r^2 = 0.95$ for the benchmark on simulated maps), with a best-fit line of $y = 0.61x + 2.4$. This linear relationship was used to predict model accuracy for all models built into experimental density maps. Per-residue convergence was similarly calculated over the top ten scoring models and also correlates with per-residue model accuracy ($r^2 = 0.88$ for the benchmark on simulated maps), with a best-fit line of $y = 0.75x + 2.0$. Real-space correlation coefficients over the full model and per residue were calculated in Rosetta with the following command (*34*):

```
density_tools -s {model} -mapfile {density map} -mapreso {map
resolution} -cryo-EM_scatterers -denstools::perres
```

hc16 mutate-map-rescue experiments

All mutate-map-rescue experiments were performed on the hc16 product construct used for the M2-seq experiments. For each base pair tested, we probed two single mutants that would disrupt the putative base pair and a double mutant that should restore base pairing (for a G-C base pair, the three mutations were G→C, C→G, and G→C and C→G; for an A-U base pair, the three mutations were A→U, U→A, and A→U and U→A; for a G-U base pair, the three mutations were G→C, U→G, and G→C and U→G), expect for mutations testing P11 and alt-P11. The specific mutations that were tested are listed in Fig. S11, Fig. S12, and Fig. S13. Chemical mapping was performed as described previously (*35*). Briefly, RNA was prepared as described above. Primers for PCR assembly of the DNA templates were designed with Primerize and are listed in Table S6 (*36*). Chemical mapping was performed in 50 mM Na-HEPES pH 8.0, 10 mM $MgCl_2$ with 1.06 mg/mL 1M7 in DMSO (all concentrations are final). Reactions were quenched and purified as described previously (*35*). cDNA was generated with SuperScript III. Reverse transcription reactions were performed at 48 °C for 40 minutes. cDNA purification was performed as described previously (*35*). cDNA was combined with GeneScan 350 ROX dye Size Standard (Life Technologies, 401735) to provide an internal control. Capillary electrophoresis was performed at ELIM Biopharmaceuticals, Inc. with an ABI 3130 or ABI 3730 machine.

### Supplementary Text

Screening RNA molecules with native gels

As the first step of the Ribosolve pipeline, RNA molecules were screened on a native gel (Fig. 1A). All of the molecules in our benchmark set ran as sharp bands in our standard buffer conditions (Methods), with the exceptions of the 24-3 ribozyme and downstream peptide riboswitch (Fig. S2). We envision that this screening step could be used to optimize folding conditions or identify molecules that may be poorly suited to structural characterization with cryo-EM. Our results suggest that it is necessary, but not sufficient for RNA molecules to pass this check.

Assessing auto-DRRAFTER accuracy with simulated density maps

To assess the accuracy of auto-DRRAFTER, we benchmarked the method using a set of eight RNA molecules with previously solved crystal structures. We used auto-DRRAFTER to build models into 10 Å resolution simulated density maps. The final models closely resembled the crystal structures, with the best RMSD accuracy of the top ten scoring auto-DRRAFTER models ranging from 2.3 – 10.0 Å (Table S2, Fig. S8A-H). Previously, we showed that DRRAFTER model accuracy for RNA coordinates built into cryo-EM maps of large RNPs could be reliably predicted from modeling convergence, which is defined as the average pairwise RMSD over the top ten scoring models (*26*). We confirmed that there is also a strong correlation between auto-DRRAFTER modeling convergence and average model RMSD accuracy ($r^2$ = 0.95), suggesting that auto-DRRAFTER model accuracy can also be reliably predicted (Fig. 1C). We also confirmed that convergence correlates with accuracy for individual residues ($r^2$ = 0.88), though there are several outliers for residues that have very low convergence, suggesting that per-residue convergence values should be interpreted with caution (Fig. S9A). Model accuracy is also correlated with real-space correlation coefficients (CC) for the map versus model (full model $r^2$ = 0.76, per residue $r^2$ = 0.48), suggesting that CC can provide an additional check of final auto-DRRAFTER model accuracy (Fig. S9B, C).

Using convergence and map vs. model CC to assess Ribosolve model accuracy

To assess the accuracy and quality of our models, we looked at the overall and per-residue convergence and CC. Based on the results from our auto-DRRAFTER benchmark on simulated maps, we expect these values to correlate with model accuracy (Fig. 1C, Fig. S9). We first confirmed that this was the case for the *F. nucleatum* glycine riboswitch, for which a crystal structure of nearly the entire riboswitch has previously been solved. For a set of generally inaccurate models (mean RMSD to the crystal structure = 14.1 Å), the overall convergence was high (20.2 Å), nearly all residues had convergence values above 10 Å, and many residues had CC values below 0.5, indicating that they did not fit well in the density map (Fig. S18A-C). In contrast, for a set of generally accurate models (mean RMSD to the crystal structure = 4.9 Å), the convergence RMSD over all residues was low (3.2 Å), nearly all individual residues had convergence values below 10 Å, and all residues had CC values above 0.5 (Fig. S18D-F).

Combining M2-seq secondary structure determination with auto-DRRAFTER

Auto-DRRAFTER requires RNA secondary structure as an input. For the tests using simulated maps and for most of our Ribosolve benchmark set, the secondary structures were known or hypothesized based on phylogenetic covariance analysis. Secondary structures were not known for the synthetic and *in vitro* evolved RNAs for which only single sequences were available in the literature, precluding covariance analysis. To elucidate the unknown secondary structures and to confirm hypothesized secondary structures, we used a multidimensional chemical mapping technique called mutate-and-map read out by next-generation sequencing (M2-seq) (*6*). M2-seq has previously been shown to be particularly accurate in revealing specific base-pairing interactions, and can now be carried out quickly and in a single pot (*6*). The M2-seq-based secondary structures for all RNAs in our benchmark set agreed well with previous models based on sequence covariation, previously solved crystal structures, or designs, where available (Fig. S5, Fig. S6, Fig. S7). These M2-seq-based secondary structures were then used as inputs along with our cryo-EM maps for the auto-DRRAFTER pipeline.

Fully automated and best-case model building for thirteen RNAs

For each of the thirteen RNAs for which we were able to obtain cryo-EM maps, we built fully automated Ribosolve models without any user intervention as well as "best-case" Ribosolve models using additional information such as placement of sub-structures within the density maps, previously solved crystal structures of parts of the RNA molecules, and modifications to the M2-seq-based secondary structures based on sequence covariation and previously solved crystal structures (see Methods, Fig. S6). The best-case models for all systems converged well (predicted RMSD accuracy < 6.3 Å, Fig. S19, Table 1) and the automated models for all systems except the *Tetrahymena* ribozyme, hc16, the hc16 product, and the SAM-IV riboswitch with SAM converged well (predicted RMSD accuracy < 4.8 Å, Fig. S20, Table S4). The M2-seq-based secondary structures used for the auto-DRRAFTER modeling for the SAM-IV riboswitch with SAM and the *Tetrahymena* ribozyme contained base pairs that were incompatible with sequence covariation for the SAM-IV riboswitch and previously solved crystal structures for the *Tetrahymena* ribozyme (Fig. S6A, L). The majority of these base pairs were automatically flagged as uncertain by the M2-seq analysis (bootstrap probabilities less than 90%). For the best-case modeling, these secondary structures were modified to resolve these inconsistencies, which may explain why the best-case models for these systems converged while the automated models did not (Fig. S6A, L). We also needed to modify the M2-seq-based secondary structures for hc16 and the hc16 product to be able to build models that converged and fit in the density maps. Again, this required removing base pairs with bootstrap probabilities less than 90. Further details are described below.

Model building and prospective compensatory mutagenesis for hc16

When the automated models for hc16 and the hc16 product initially did not converge, we hypothesized that like for the SAM-IV riboswitch with SAM and the *Tetrahymena* ribozyme, the secondary structure required refinement. Notably, hc16 was evolved *in vitro* from a random library that contained the P4-P6 domain of the *Tetrahymena* ribozyme as a constant scaffold region (*4*), but the M2-seq-based secondary structure was inconsistent with the presence of this structural element. In particular, the M2-seq data suggested that the P4 stem from the *Tetrahymena* ribozyme was replaced by an alternative helix, which we will refer to as alt-P4 (Fig. S21A, B). We instead tried to build models using the previously hypothesized secondary structure, which contains the P4 stem from the *Tetrahymena* ribozyme, but the models still did not fit well in the density maps. To resolve whether P4 or alt-P4 was formed in hc16 and the hc16 product, we performed mutate-map-rescue experiments (*35*). In addition to P4 and alt-P4, we also tested P7 for which we could not see a clear signal in the M2-seq data and a possible alternative, alt-P7. Our results support the formation of the alt-P4 helix observed in the M2-seq data as well as P7, and do not provide evidence for the formation of P4 or alt-P7 (Fig. S11).

New sets of auto-DRRAFTER models with an updated secondary structure containing alt-P4, P7, and additional previously hypothesized secondary structure elements that were not present in the M2-seq-based secondary structure (Fig. S21B) still did not converge and did not fit well in the density maps. Again, we reexamined the secondary structure and, based on the discrepancies between our models and the density maps and regions marked uncertain by M2-seq secondary structure analysis, we hypothesized several modifications to the secondary structure: an extension of the alt-P4 helix that would replace P5, possible additional helices (P10, P12), and possible two base pair pseudoknots (P11/alt-P11) (Fig. S21C). Additional mutate-map-rescue experiments support the replacement of P5 with alt-P4, and provide some evidence for P10, but do not clearly discriminate between P11 and alt-P11 and P10-ext and P12 (Fig. S12 and Fig. S13). We therefore built auto-DRRAFTER models with each possible secondary structure. The final ensemble of auto-DRRAFTER models reflects this uncertainty in secondary structure, but still converges well (6.2 Å for hc16, 6.3 Å for hc16 product, and 5.5 Å for the second conformation of the hc16 product) and fits well in the density map (CC = 0.81 for hc16, CC = 0.79 for hc16 product, CC = 0.79 for the second conformation of the hc16 product).

Evaluating the accuracy of Ribosolve models

Our auto-DRRAFTER benchmark suggested that overall and per-residue convergence and correlation between the map and model (CC) would correlate with model accuracy (Supplementary Text, Fig. S9, Fig. 1C). Therefore, as an initial check of the accuracy of the Ribosolve models, we confirmed that the per-residue convergence values were generally below 10 Å and the per-residue and overall CC values were above 0.5 for all automated and best-case models that converged (overall convergence < 10 Å, Fig. S20, Fig. S19). Residues that do not fit these criteria are listed in Table S5.

The *F. nucleatum* and *V. cholerae* glycine riboswitch Ribosolve models agreed well with the previously solved crystal structures (*30, 33*) with mean RMSDs over the top ten scoring models refined into each half map of 4.9 Å and 8.6 Å for the *F. nucleatum* glycine riboswitch with and without glycine, respectively, and 3.3 Å and 3.6 Å for the *V. cholerae* glycine riboswitch with and without glycine, respectively (Fig. S10A, B). For the *Tetrahymena* ribozyme, best-case models were built starting from the previously solved crystal structure of the core of the ribozyme. The complete *Tetrahymena* ribozyme Ribosolve models fit well in the density map, with little deviation from the crystal structure in the core (RMSD = 3.5 Å, Fig. S10C, D).

For hc16 and the hc16 product, we confirmed that the Ribosolve models are consistent with all previous observations of the ribozyme (*4*). First, our models contain the P7 and P9 helices, which each contain nucleotides that exhibited covariation in clones examined from the original *in vitro* selection experiments. Nucleotides that were invariant across the final *in vitro* selected sequences are shown as red spheres in Fig. S10E and are all automatically modeled as near the substrate binding site or forming base pairs with regions of fixed sequence in the selection libraries. Nucleotides that were not conserved between sequences are shown as white spheres in Fig. S10E and do not make any interactions with other parts of the ribozyme in our model. Additionally, removing the thirteen nucleotides at the 3' end of the ribozyme abolishes its activity. In our model, most of these nucleotides are part of the P10, P10-ext, or P12 stems in the core of the structure.

The M2-seq data for the ATP-TTR-3 with and without AMP, spinach-TTR-3, and eterna3D-JR_1 molecules contain features that are not explained by the M2-seq-based secondary structures and appear to correspond to tertiary contacts, providing additional information about nucleotides that should be close together in the three-dimensional structures (Fig. S10F, Fig. S14). We confirmed that our Ribosolve models, which were built without using this information, recover these tertiary contacts (Fig. S10G, Fig. S14).

Finally, six nucleotides in the SAM-IV riboswitch were previously hypothesized to form the binding pocket for SAM (*37*). We confirmed that these nucleotides are all located near each other in the fully automated SAM-IV apo state Ribosolve models, which were not built using this information, and that the conformation of these nucleotides is in good agreement with the conformation of putatively homologous nucleotides in the previously solved crystal structure of the SAM-I riboswitch (RMSD = 3.3 Å, Fig. S10H). Additionally, the best-case models for SAM-IV with and without SAM, which were built using this homology (see Methods for details), agree well with the fully automated models (RMSD = 5.0 Å and 5.6 Å for best-case models with and without SAM, respectively, vs. fully automated models without SAM).

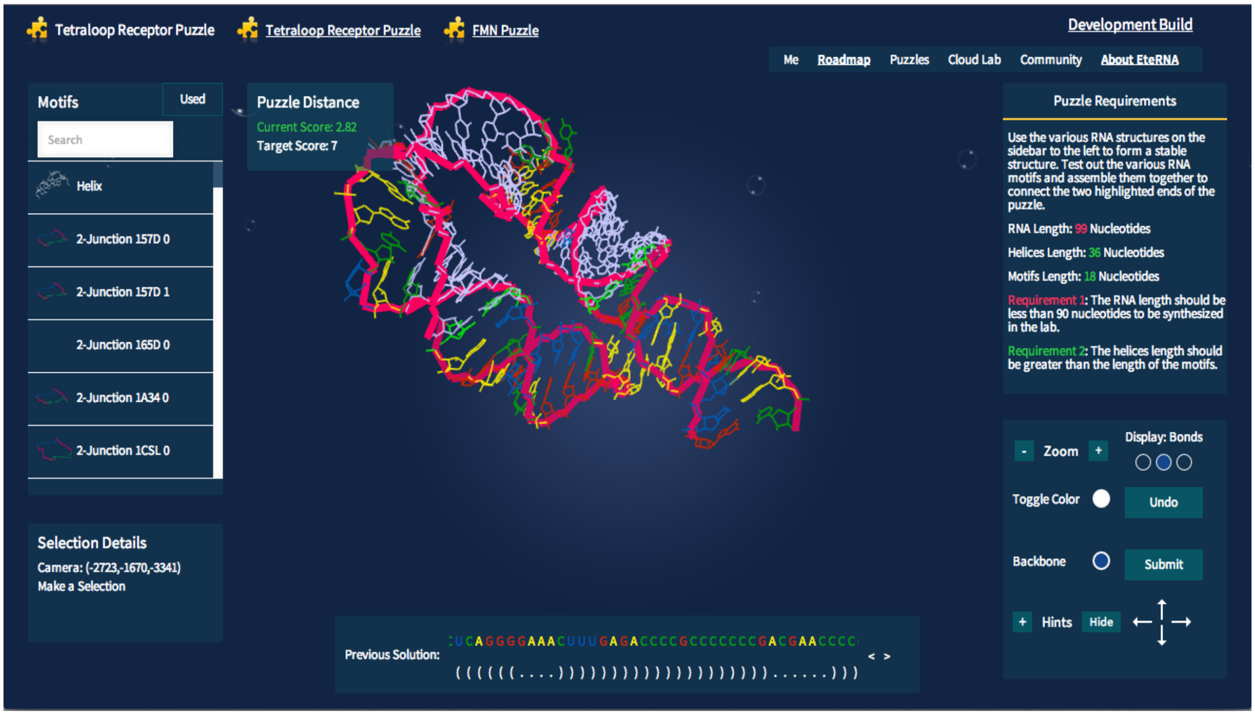


**Fig. S1.**
The interface used by Eterna players to rationally design RNA segments that connect the two segments of the tetraloop/tetraloop-receptor contact. Players were able to utilize both two-way junction motifs and helices from a list on the left to build a path from the tetraloop to the tetraloop receptor.

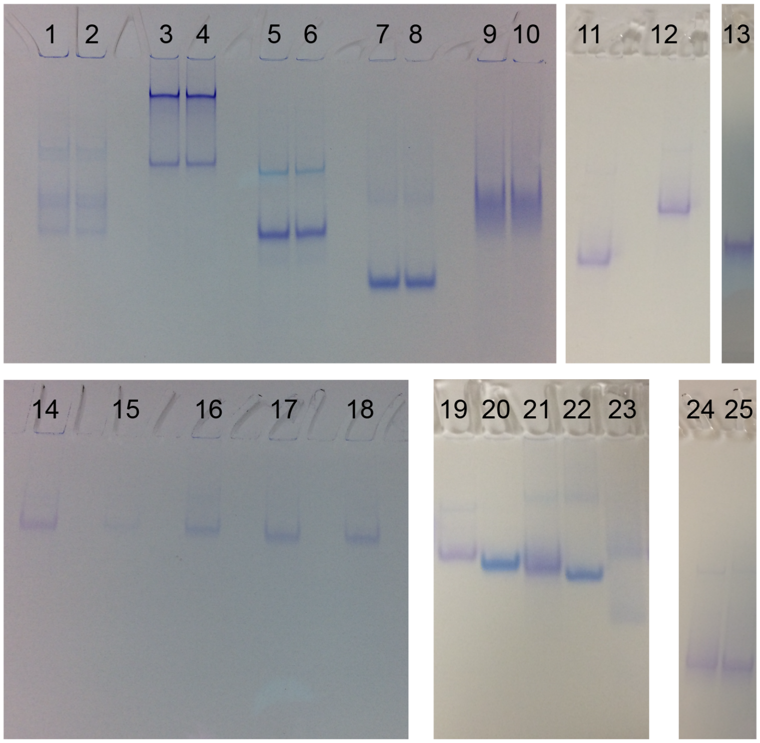


**Fig. S2.**

Native gels for (1) and (2) RB1 5′ UTR, (3) and (4) scaRNA6, (5) and (6) U1 snRNA, (7) and (8) SAM-IV riboswitch (apo), (9) and (10) 24-3, (11) *V. cholerae* glycine riboswitch (apo), (12) hc16, (13) *Tetrahymena* ribozyme, (14) *F. nucleatum* glycine riboswitch (apo), (15) and (16) Eterna3D-JR_1, (17) and (18) spinach-TTR-3, (19) *F. nucleatum* glycine riboswitch (apo), (20) ATP-TTR-3 (apo), (21) Eterna3D-JR_1, (22) SAM-IV riboswitch (apo), (23) downstream peptide riboswitch, (24) hc16, and (25) hc16 product. Samples 1-13, 24, 25 were run on an 8% polyacrylamide gel. Samples 14-23 were run on a 12% polyacrylamide gel. All samples were run in 10 mM $MgCl_2$, 67 mM HEPES, 33 mM Tris, pH 7.2.x

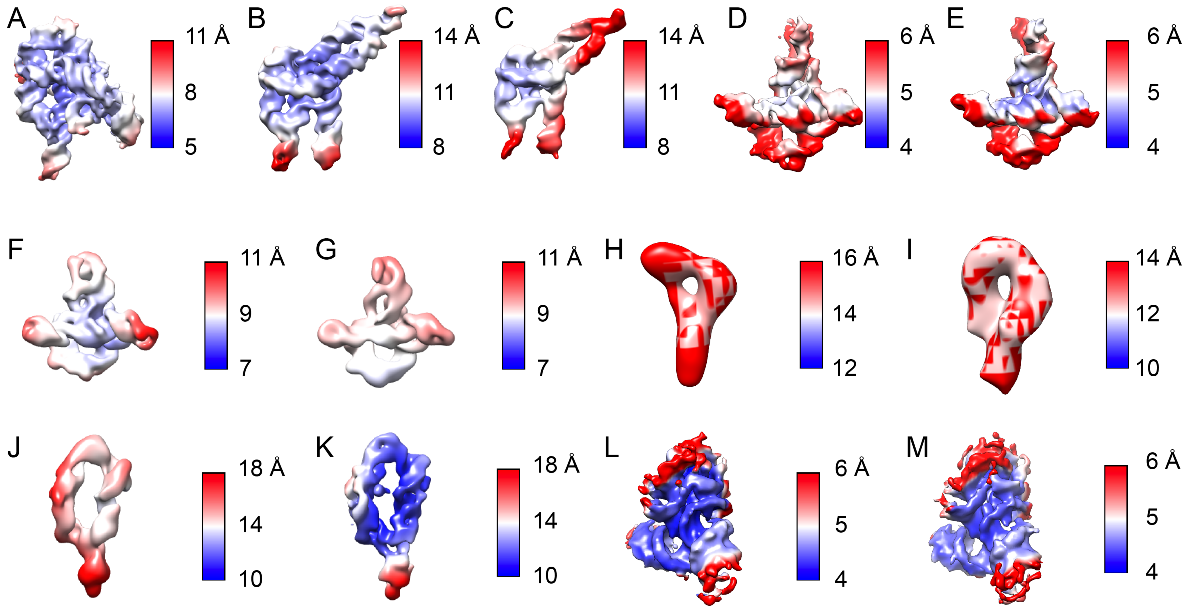


**Fig. S3.**

Local map resolution, calculated with ResMap (*38*) for (A) *Tetrahymena* ribozyme, (B) hc16 product, (C) hc16, (D) *V. cholerae* glycine riboswitch with glycine, (E) *V. cholerae* glycine riboswitch without glycine, (F) *F. nucleatum* glycine riboswitch with glycine, (G) *F. nucleatum* glycine riboswitch without glycine, (H) eterna3D-JR_1, (I) spinach-TTR-3, (J) ATP-TTR-3 with AMP, (K) ATP-TTR-3 without AMP, (L) SAM-IV riboswitch with SAM, and (M) SAM-IV riboswitch without SAM.

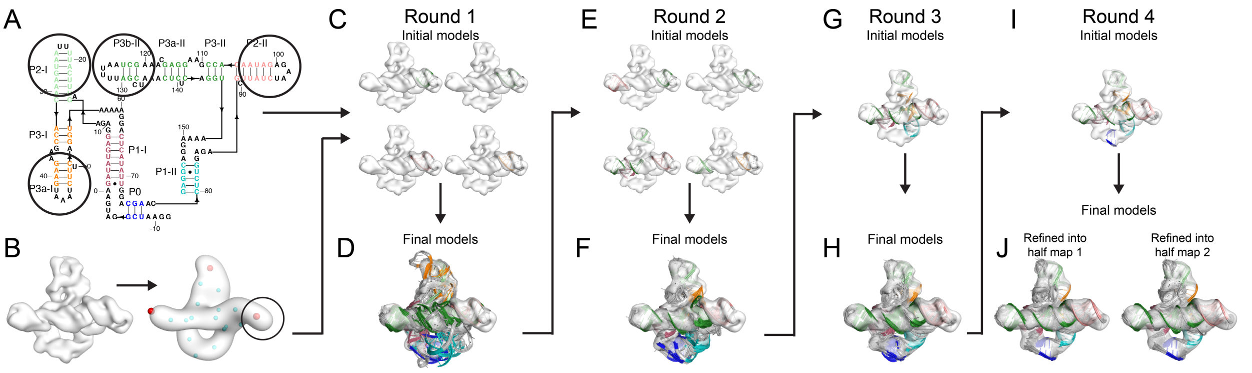


**Fig. S4.**

auto-DRRAFTER overview. The *F. nucleatum* glycine riboswitch is shown here as an example. (A) Secondary structure elements that connect to just one other helix or junction ("end nodes") are circled. (B) The cryo-EM density map is low-pass filtered to 20 Å and points are placed (spheres) throughout the map to identify possible placements for "end nodes" in the map (red spheres). The circled end node was randomly selected for initial helix placement. (C) 3D models are built for each of the elements circled in (A) and fit into the density map in the location of the circled point in (B). These elements are kept fixed while the rest of the RNA is built into the density map. (D) The top ten best scoring models after round 1. The overall convergence of these models is above the 10 Å threshold (convergence = 20.2 Å), so another round of modeling is performed. (E) For the second round of modeling, regions that have converged are extracted from the top scoring models and kept fixed while the rest of the RNA is built into the density map. (F) The top ten scoring models after round 2. The convergence is below the 10 Å threshold (convergence = 6.2 Å), so there is only one initial model for the third round of modeling, composed of converged regions from the top ten scoring models from round 2 (G). These regions are allowed to move from their initial positions during this modeling round. (H) The best scoring models from round 3. (I) Again, converged regions are extracted from top scoring models to form the initial model for the final round of modeling. These regions are kept fixed during the fragment assembly stage of auto-DRRAFTER modeling, but allowed to move during final refinement. (J) The top ten scoring models built independently into each half map.

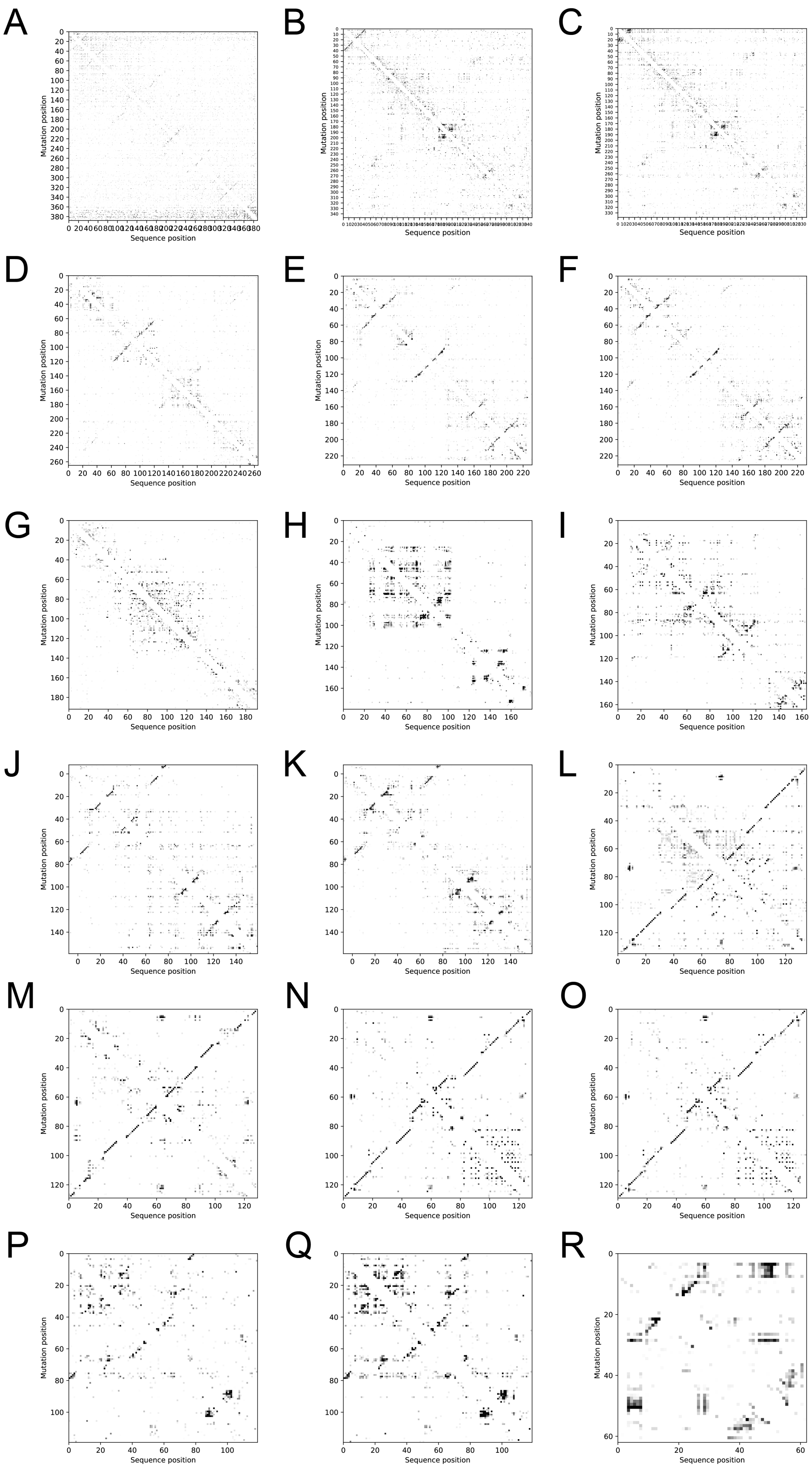
A
B
C
D
E
F
G
H
I
J
K
L
M
N
O
P
Q
R
Mutation position
Sequence position

**Fig. S5.**

Experimental M2-seq Z-score plots for (A) *Tetrahymena* ribozyme, (B) hc16 product, (C) hc16, (D) human scaRNA6, (E) *V. cholerae* glycine riboswitch with glycine, (F) *V. cholerae* glycine riboswitch without glycine, (G) human RB1 5′ UTR, (H) 24-3, (I) human U1 snRNA, (J) *F. nucleatum* glycine riboswitch with glycine, (K) *F. nucleatum* glycine riboswitch without glycine, (L) eterna3D-JR_1, (M) spinach-TTR-3, (N) ATP-TTR-3 with AMP, (O) ATP-TTR-3 without AMP, (P) SAM IV riboswitch with SAM, (Q) SAM IV riboswitch without SAM, and (R) downstream peptide riboswitch.

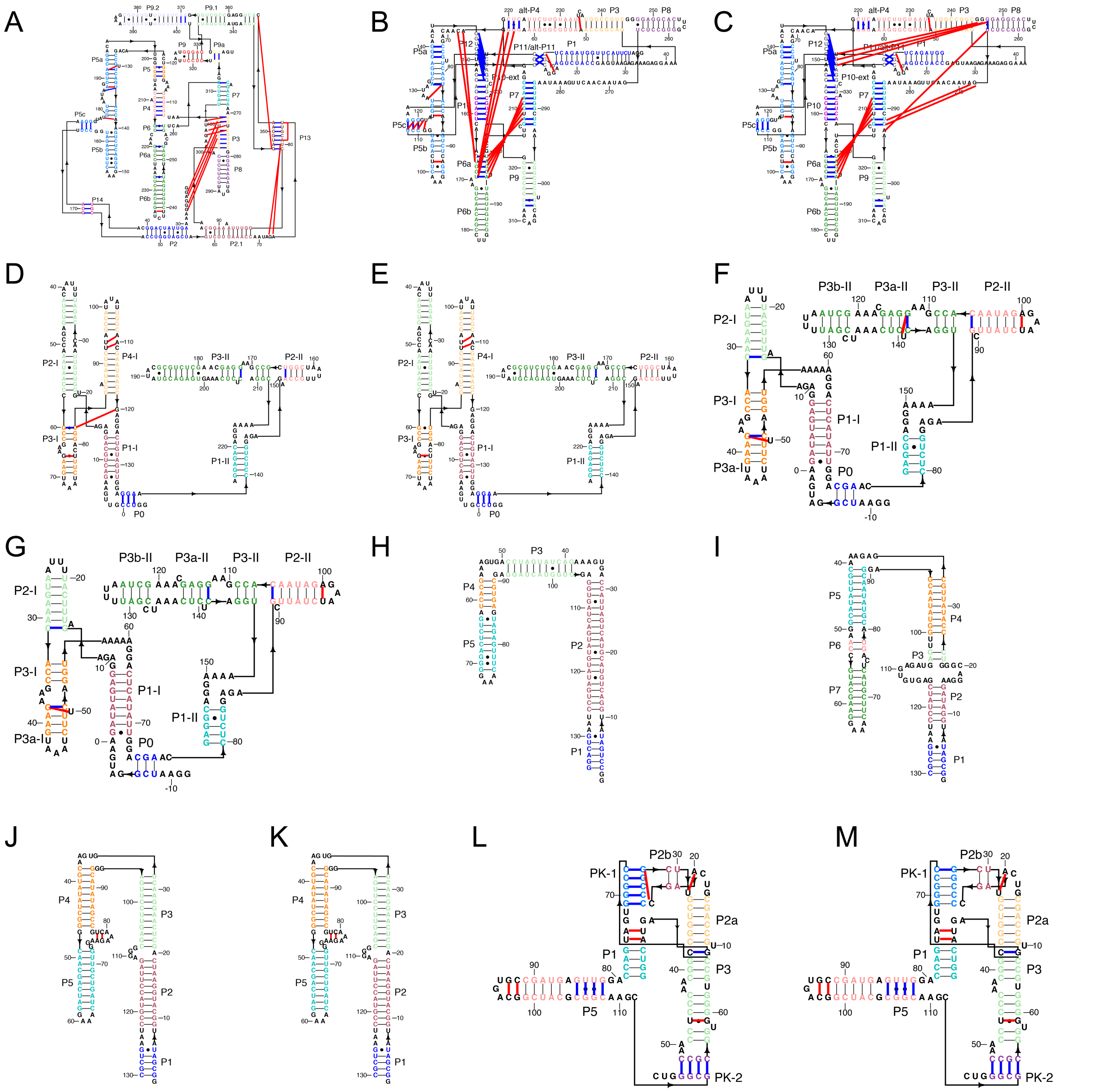


**Fig. S6.**

Comparison between the secondary structures automatically derived from M2-seq data (used for automated auto-DRRAFTER modeling) and the secondary structures used for best-case auto-DRRAFTER modeling for (A) *Tetrahymena* ribozyme, (B) hc16 product, (C) hc16, (D) *V. cholerae* glycine riboswitch with glycine, (E) *V. cholerae* glycine riboswitch without glycine, (F) *F. nucleatum* glycine riboswitch with glycine, (G) *F. nucleatum* glycine riboswitch without glycine, (H) eterna3D-JR_1, (I) spinach-TTR-3, (J) ATP-TTR-3 with AMP (K) ATP-TTR-3 without AMP, (L) SAM-IV riboswitch with SAM, and (M) SAM-IV riboswitch without SAM. Blue lines indicate base pairs that are present in the best-case, but not automated secondary structures. Red lines indicate base pairs that are present in the automated, but not best-case secondary structures.

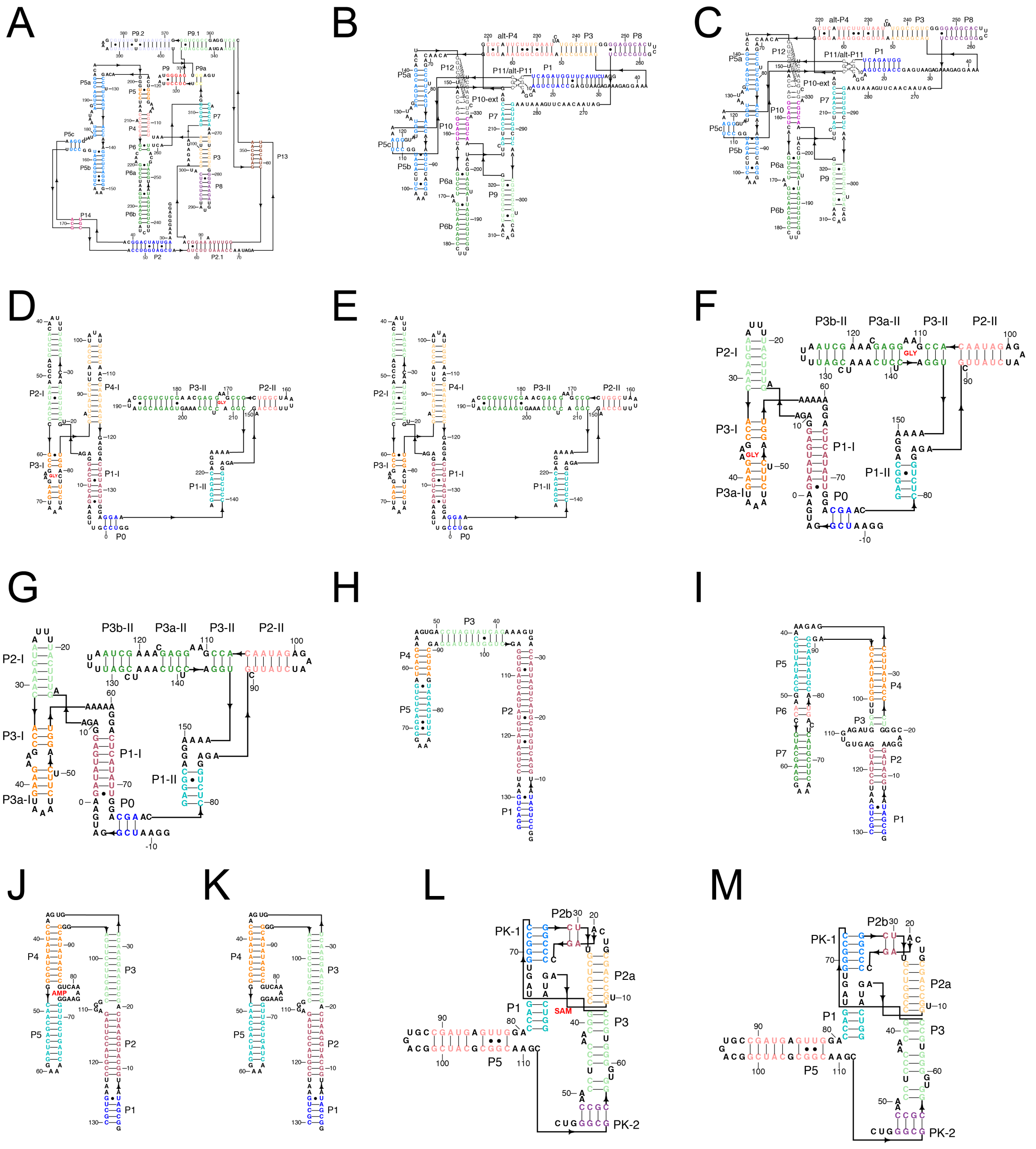


**Fig. S7.**

Secondary structures for best-case Ribosolve models for (A) *Tetrahymena* ribozyme, (B) hc16 product, (C) hc16, (D) *V. cholerae* glycine riboswitch with glycine, (E) *V. cholerae* glycine riboswitch without glycine, (F) *F. nucleatum* glycine riboswitch with glycine, (G) *F. nucleatum* glycine riboswitch without glycine, (H) eterna3D-JR_1, (I) spinach-TTR-3, (J) ATP-TTR-3 with

AMP (K) ATP-TTR-3 without AMP, (L) SAM-IV riboswitch with SAM, and (M) SAM-IV riboswitch without SAM.

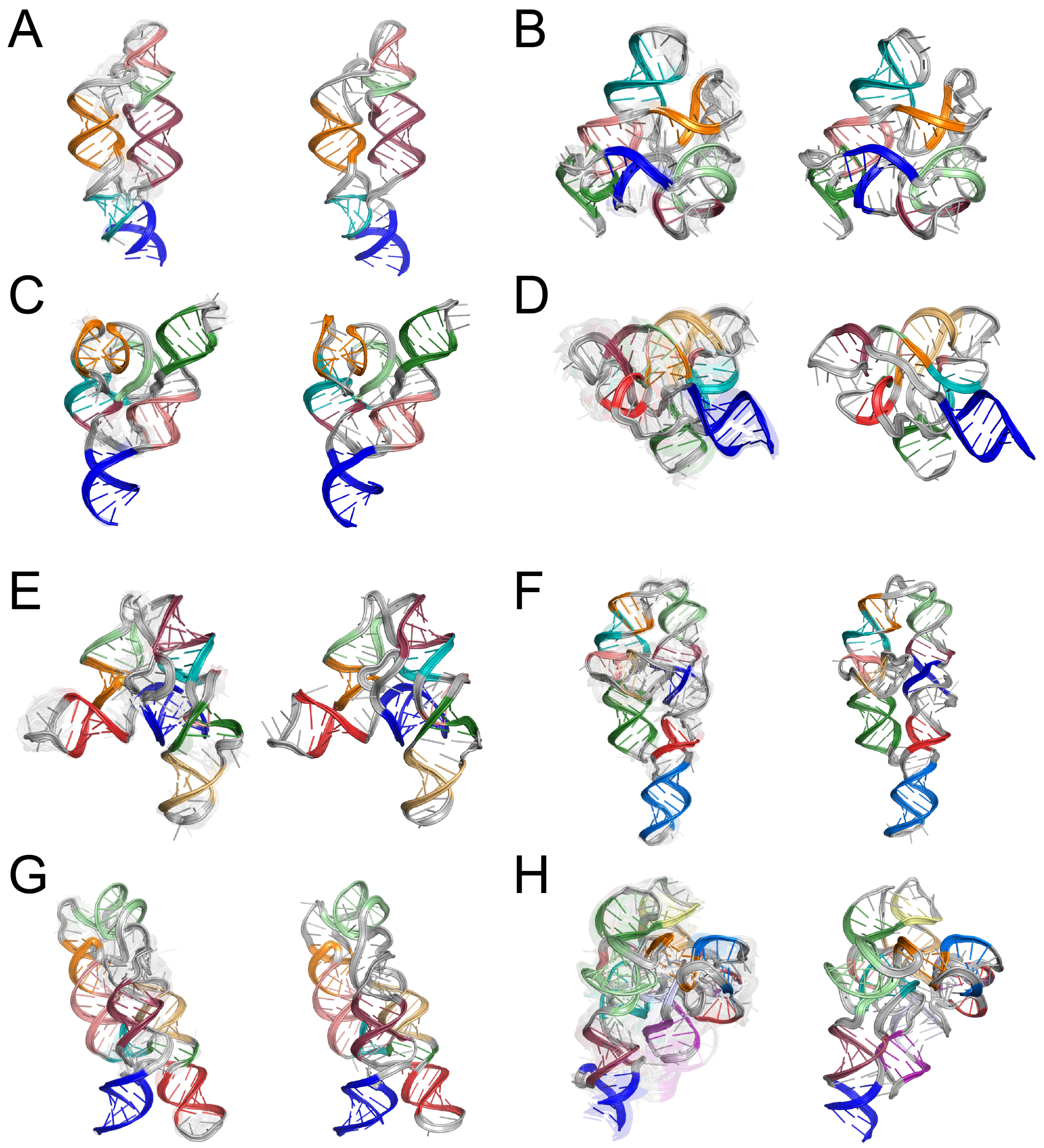


**Fig. S8.**

Benchmarking auto-DRRAFTER accuracy with simulated density maps. (A-H) The top ten scoring auto-DRRAFTER models (models #2-10 are transparent) built into 10 Å simulated density maps (left) and the corresponding crystal structures (right) for (A) THF riboswitch, (B) c-di-AMP riboswitch, (C) bacterial SRP Alu domain, (D) FMN riboswitch, (E) SAM-I riboswitch, (F) *Tetrahymena* ribozyme P4-P6 domain, (G) lysine riboswitch, and (H) lariat capping ribozyme.

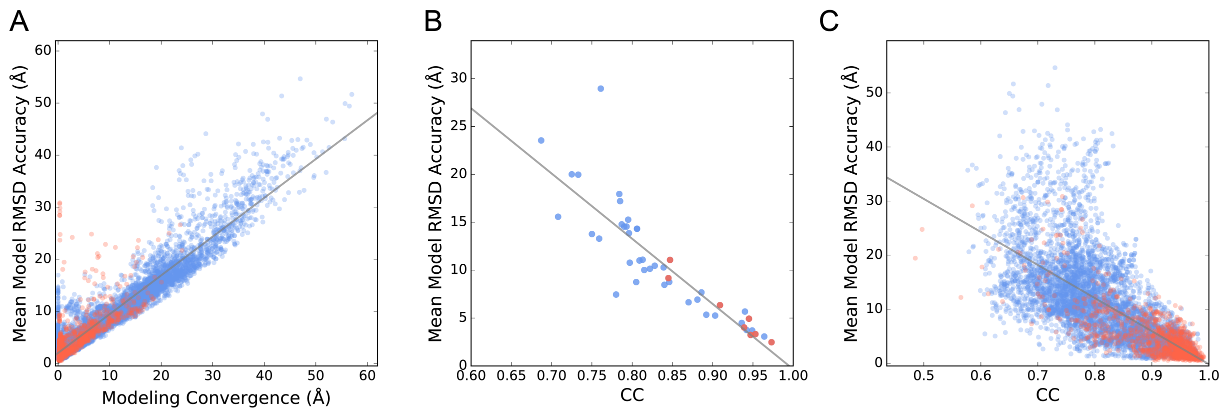


**Fig. S9.**

RMSD accuracy versus (A) convergence or (B and C) real-space CC for the top ten scoring auto-DRRAFTER models built into simulated density maps. (B) RMSDs and CC or convergence values calculated over the full models and (A and C) per residue. Points for models from the final round of auto-DRRAFTER modeling are colored red. Best-fit lines are colored gray and given by (A) y = 0.75x + 2.0, (B) y = -68.1x + 67.7, and (C) y = -60.2x + 61.0.

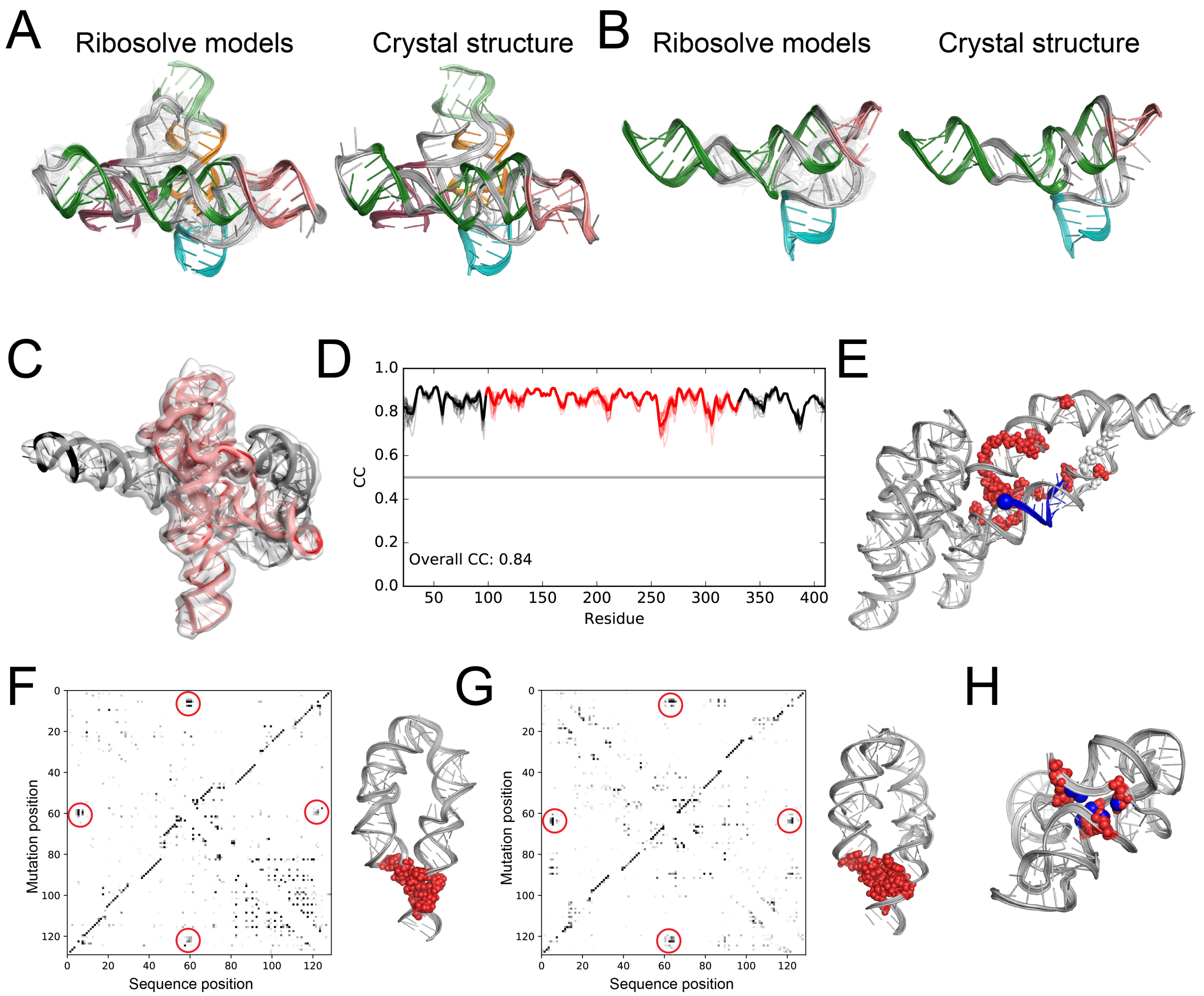


**Fig. S10.**

Independent tests of Ribosolve model accuracy. Top ten scoring fully automated Ribosolve models (models #2-10 are transparent; left) and previously solved crystal structures (right) for (A) *F. nucleatum* glycine riboswitch and (B) *V. cholerae* glycine riboswitch. Note that auto-DRRAFTER models were built for the complete RNAs, but for clarity only the parts that correspond to the previously solved crystal structures are shown. (C) The *Tetrahymena* ribozyme Ribosolve model in the cryo-EM map. The region derived from a previously solved crystal structure is colored red and regions built *de novo* are colored black. (D) Real-space correlation between the *Tetrahymena* ribozyme Ribosolve model and cryo-EM map. Colors are the same as in (C). (E) The hc16 product Ribosolve model. Invariant and highly variable residues across final sequences from the original *in vitro* selection (*4*) shown as red and white spheres, respectively. The site of substrate ligation, the phosphorus atom in residue 1, is shown as a large blue sphere. (F) and (G) M2-seq Z-score plots (left) with putative tertiary contacts circled in red and Ribosolve models with nucleotides corresponding to the circled regions in the Z-score plots shown as red spheres for (F) the ATP-TTR-3 with AMP (G) spinach-TTR-3. (H) The fully automated SAM-IV apo state Ribosolve model with nucleotides hypothesized to form the SAM binding pocket shown as red spheres. Homologous nucleotides taken from a previously solved SAM-I riboswitch structure (PDB ID: 2GIS) are overlaid and shown as blue spheres.

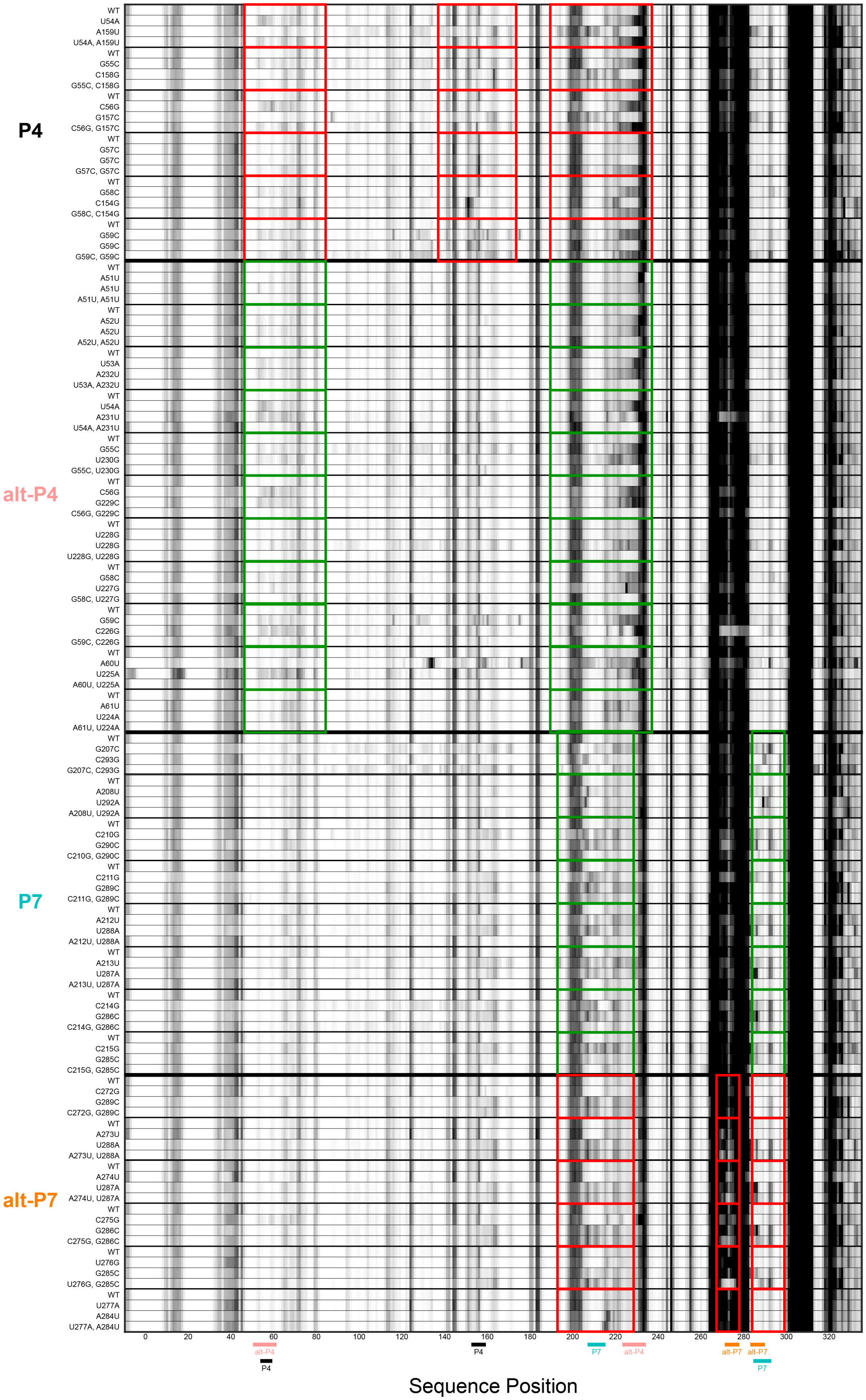

P4
alt-P4
P7
alt-P7
Sequence Position

**Fig. S11.**

Mutate-map-rescue data testing P4, alt-P4, P7, and alt-P7 in the hc16-product. Boxes denote regions where perturbations are observed. Green boxes indicate regions where perturbations are observed in the single mutants and rescue is clearly observed for the double mutants. Red boxes indicate regions where clear rescue is not observed.

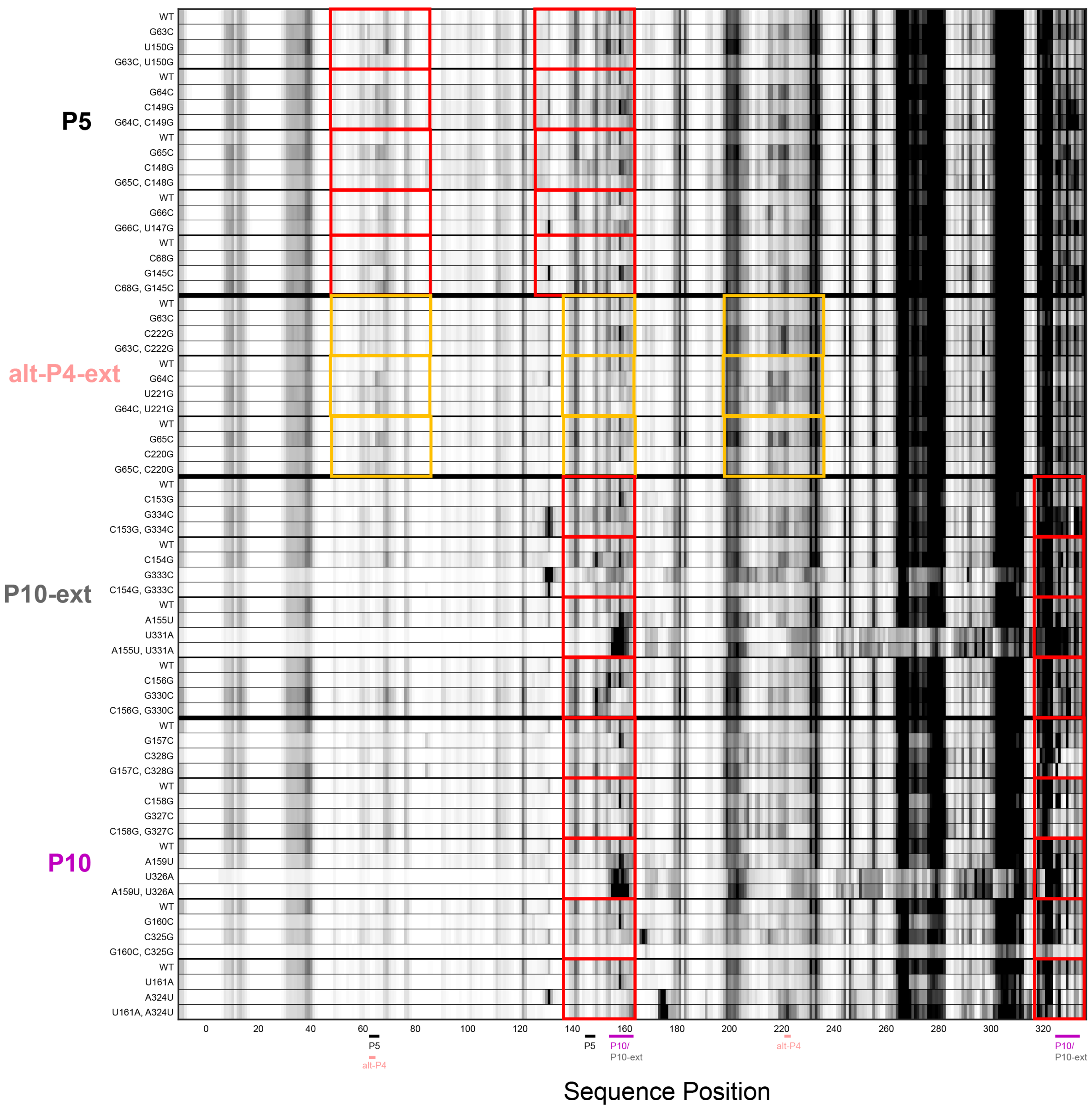


**Fig. S12.**

Mutate-map-rescue data testing P5, an extension of alt-P4, and P10 in the hc16-product. Boxes denote regions where perturbations are observed. Red boxes indicate that clear rescue is not observed. Yellow boxes indicate partial rescue.

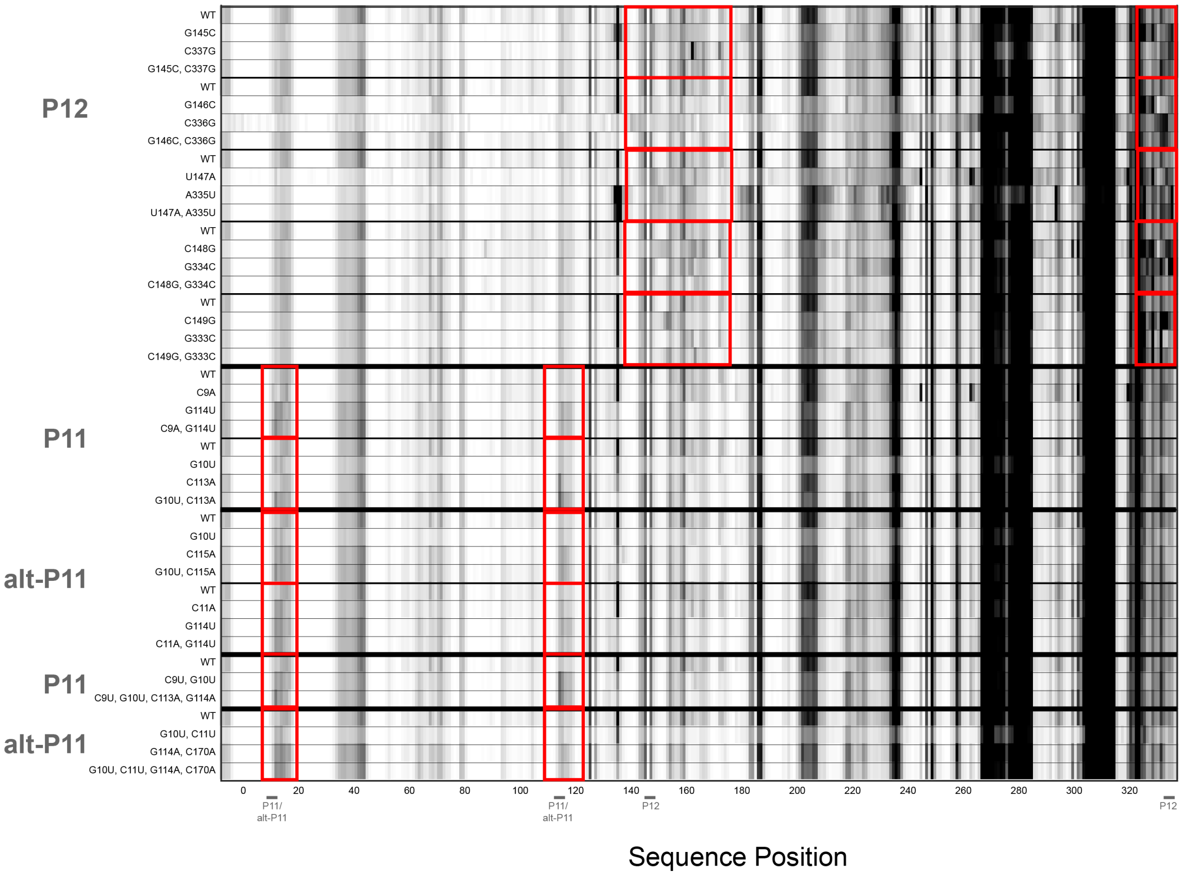


**Fig. S13.**

Mutate-map-rescue data testing P11, alt-P11, and P12 in the hc16-product. Red boxes denote regions where perturbations are observed. There is no clear rescue observed for base pairs shown here.

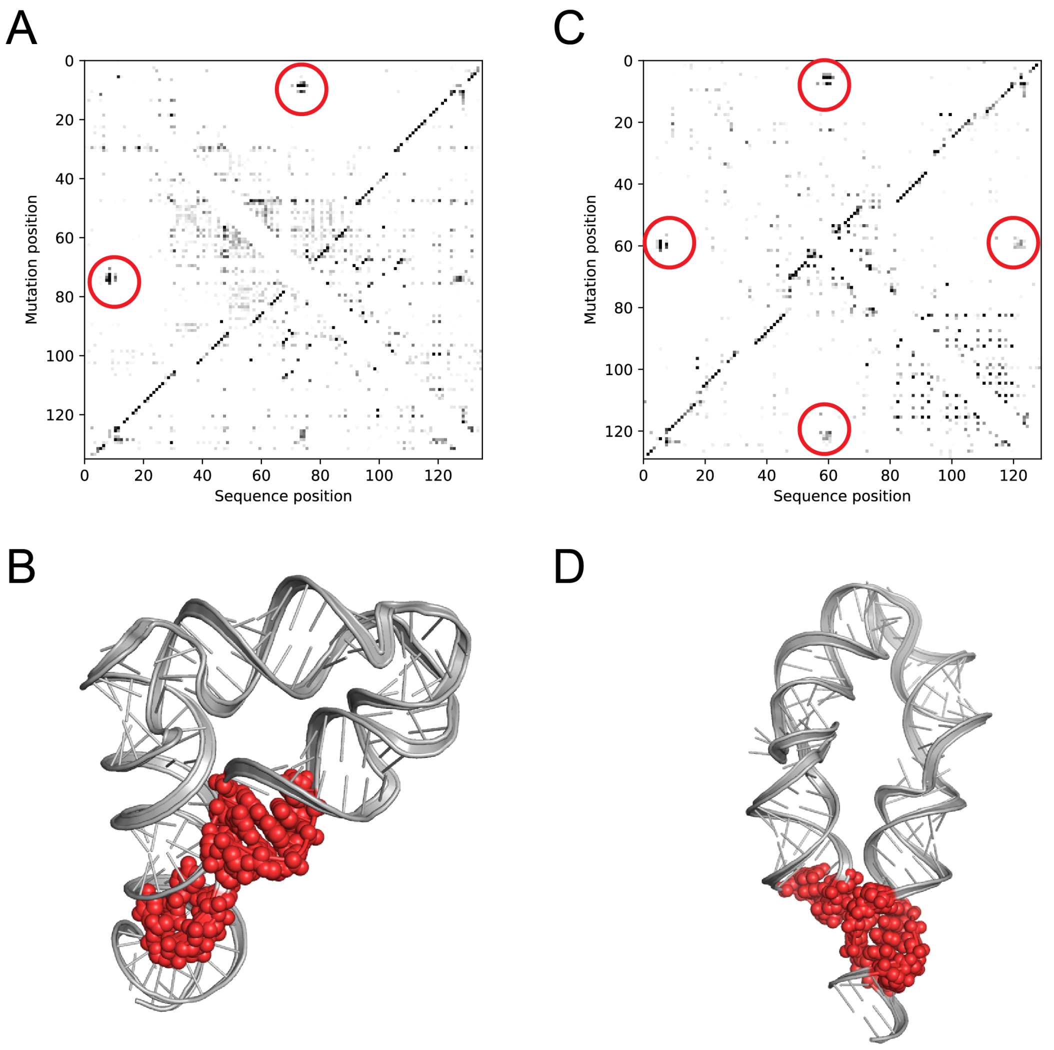


**Fig. S14.**

(A, C) M2-seq Z-score plots with putative tertiary contacts marked with red circles for (A) eterna3D-JR_1 and (C) the ATP-TTR-3 without AMP. (B, D) Ribosolve models with nucleotides corresponding to circled regions in (A) and (C) shown as red spheres for (B) eterna3D-JR_1 and (D) the ATP-TTR-3 without AMP.

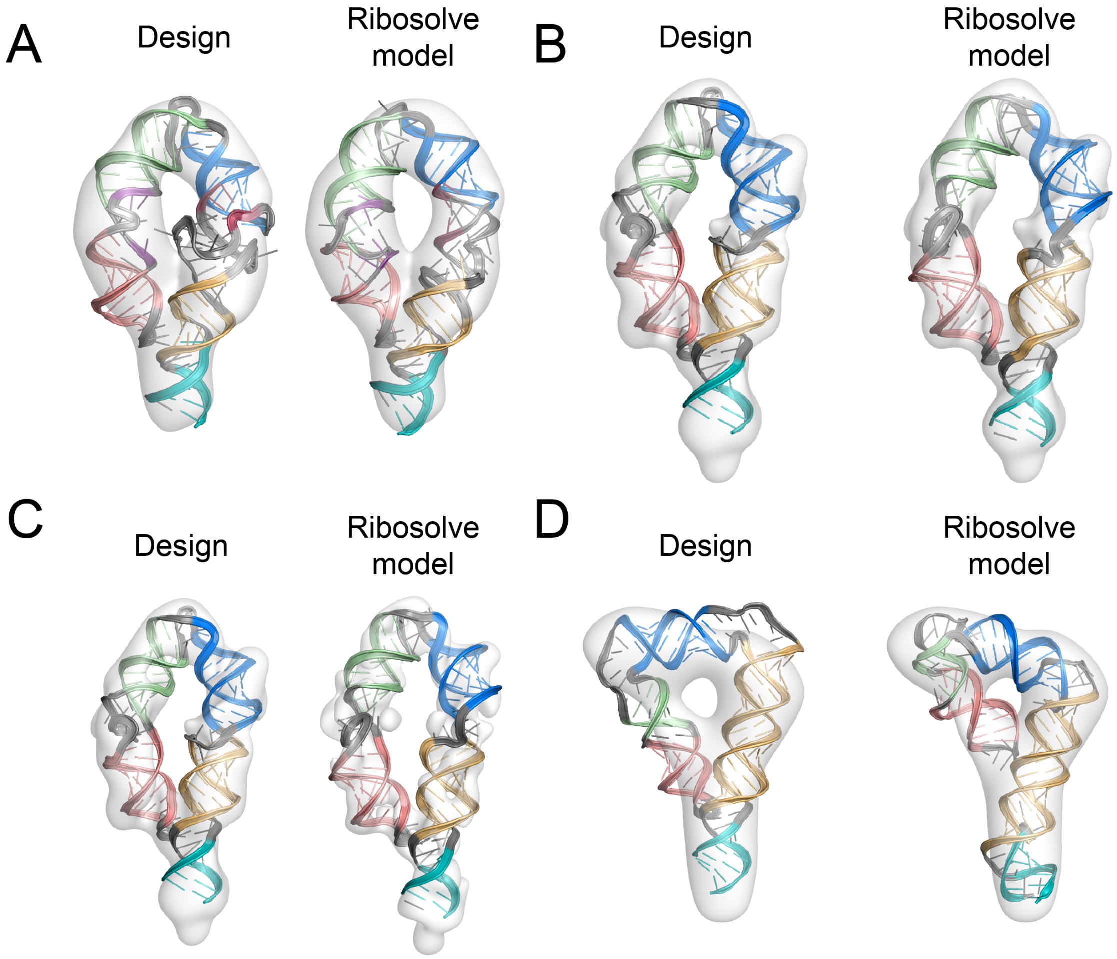


**Fig. S15.**

Comparison between computationally designed RNA structures and Ribosolve models. The computationally designed structures (left) and auto-DRRAFTER models (right) for (A) spinach-TTR-3, (B) the ATP-TTR-3 with AMP, (C) ATP-TTR-3 without AMP, and (D) Eterna3D-JR_1. Cryo-EM maps are colored gray.

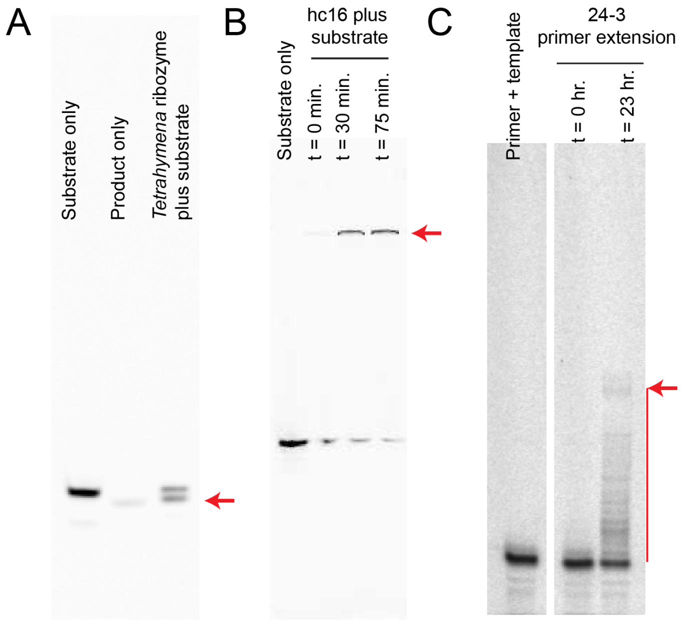


**Fig. S16.**

Ribozyme activity assays. (A) *Tetrahymena* ribozyme substrate cleavage (polyacrylamide gel, imaging Cy5, see Methods for details). (B) hc16 ligation reaction (polyacrylamide gel, imaging Cy5, see Methods for details). (C) 24-3 primer extension (polyacrylamide gel, imaging FAM, see Methods for details). Red arrows mark product(s) of ribozyme reactions.

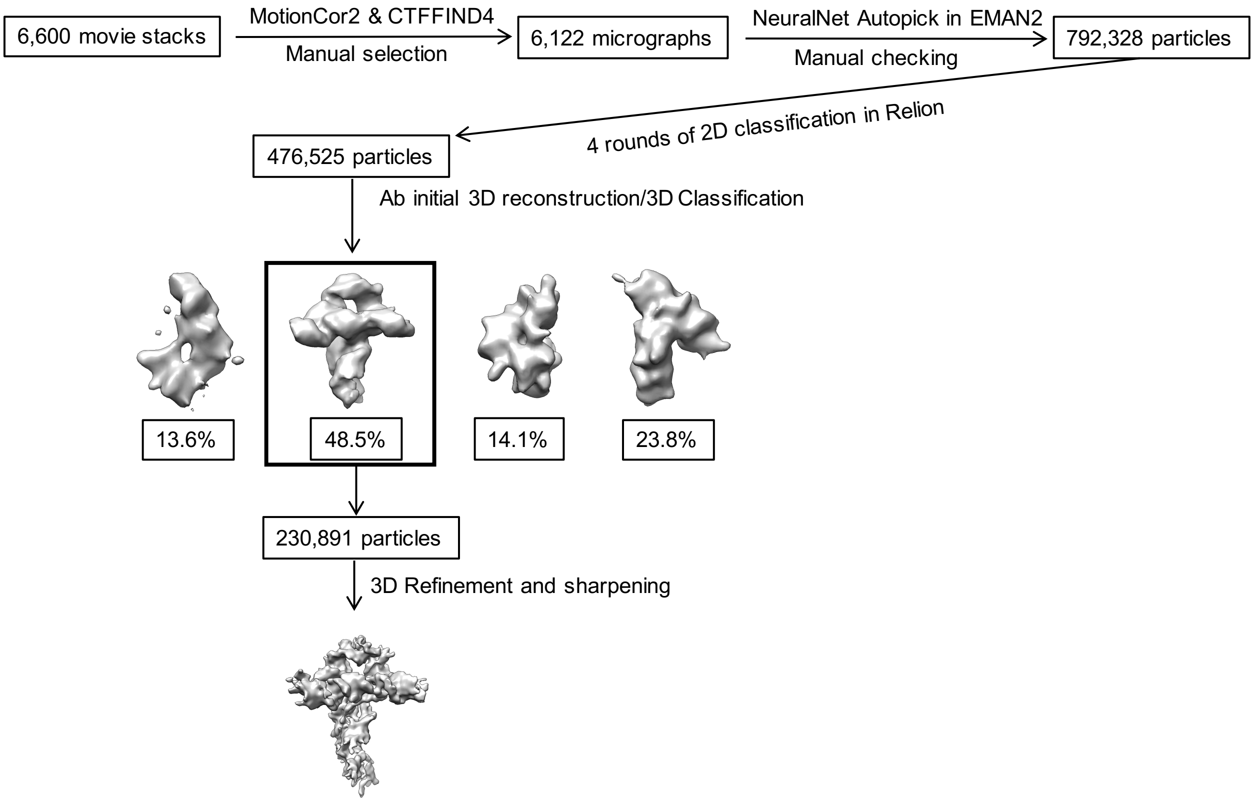


**Fig. S17.**
Example cryo-EM data processing workflow for the *V. cholerae* glycine riboswitch without glycine.

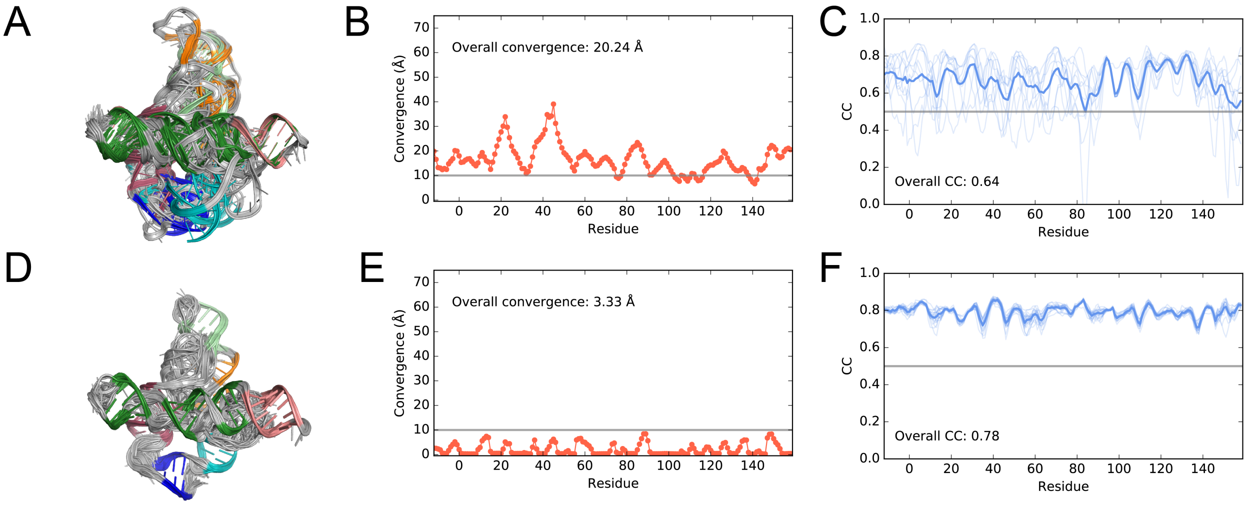


**Fig. S18.**

(A) Poorly converged and (D) well-converged auto-DRRAFTER models for the *F. nucleatum* glycine riboswitch with glycine. (B) and (E) Modeling convergence per residue for models in (A) and (D), respectively. (C) and (F) Real-space correlation coefficients (CC) per residue for the top ten scoring auto-DRRAFTER models (thin, light blue lines) for models in (A) and (D), respectively. Average CC values shown as thick blue lines.

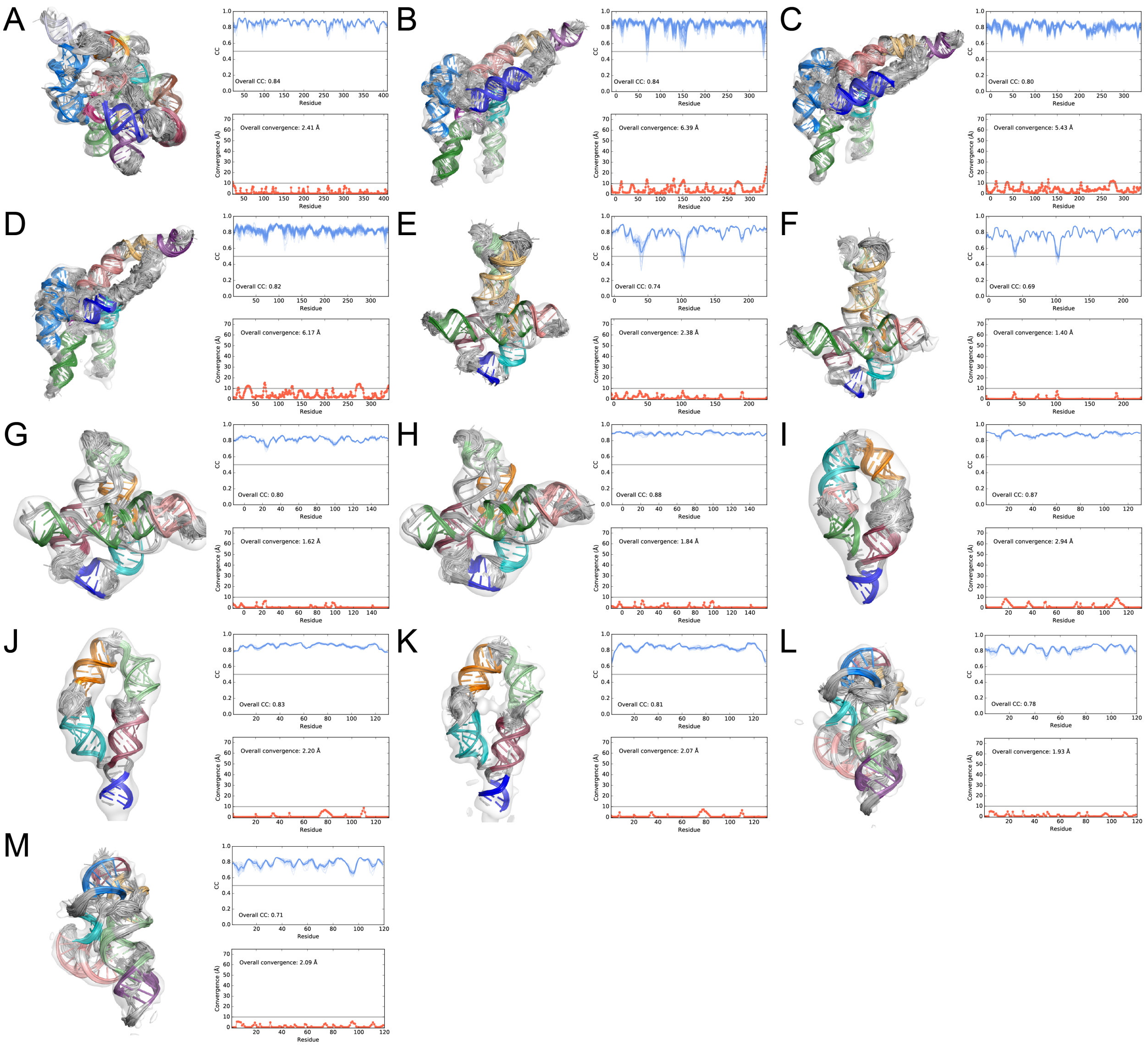


**Fig. S19.**

Best-case auto-DRRAFTER models built into experimental density maps. (A-M) (left) Top ten scoring automated auto-DRRAFTER models built into each half map shown within the full density map and (top right) the CC values between these models and the opposite half maps (i.e. models built into half map 1 are checked against half map 2). CC values for each model are shown as light blue lines, the average over all top scoring models is plotted as a thick blue line. The overall CC values reported are averaged over the top ten scoring models for each half map. (Bottom right) auto-DRRAFTER modeling convergence computed between models refined into separate half maps. (A) *Tetrahymena* ribozyme, (B) hc16 product conformation 1, (C) hc16 product conformation 2 (D) hc16, (E) *V. cholerae* glycine riboswitch with glycine, (F) *V. cholerae* glycine riboswitch, (G) *F. nucleatum* glycine riboswitch with glycine, (H) *F. nucleatum* glycine riboswitch, (I) spinach-TTR-3, (J) ATP-TTR-3 with AMP, (K) ATP-TTR-3, (L) SAM-IV riboswitch with SAM, and (M) SAM-IV riboswitch.

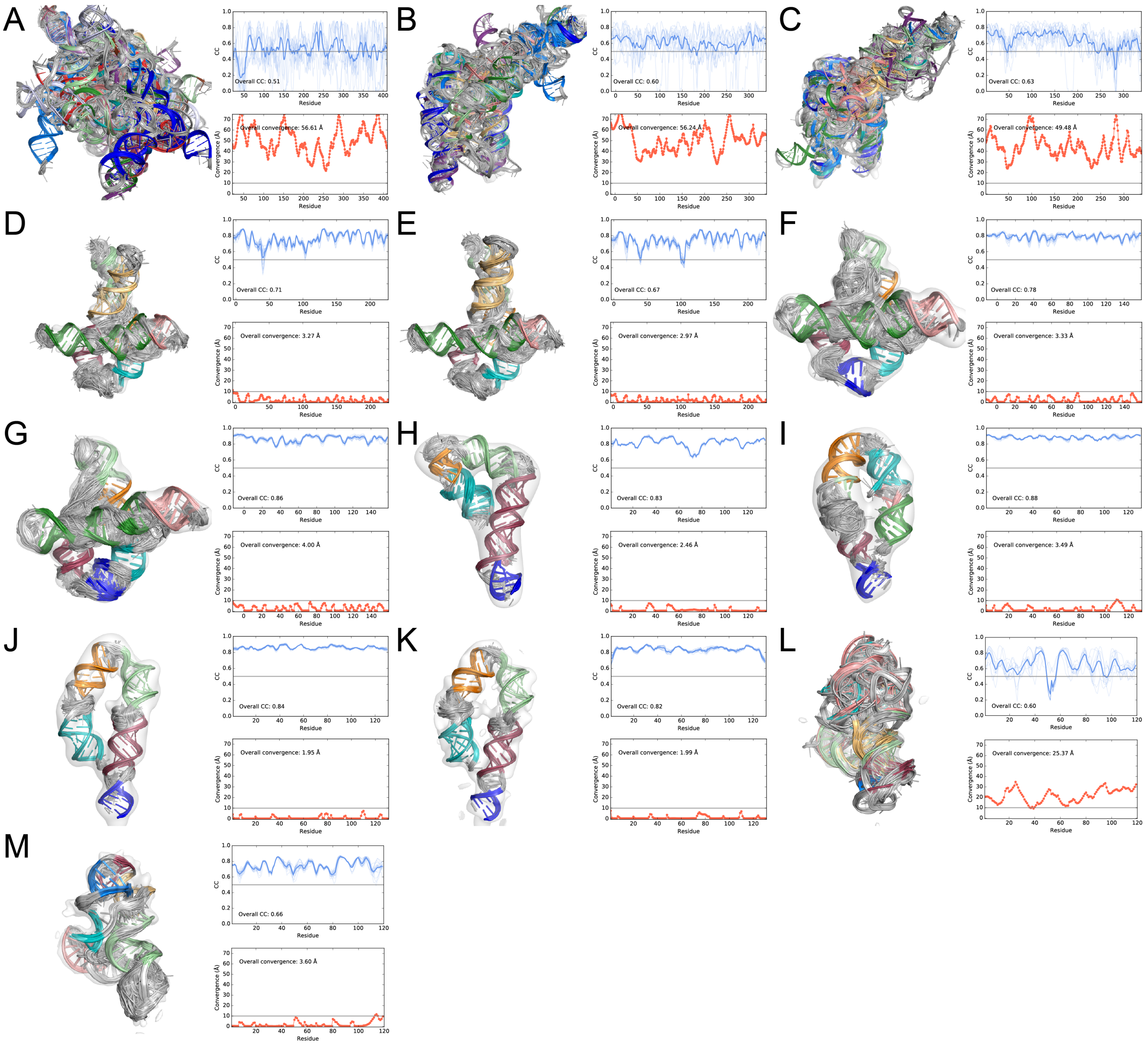


**Fig. S20.**

Automated auto-DRRAFTER models built into experimental density maps. (A-M) (left) Top ten scoring automated auto-DRRAFTER models built into each half map shown within the full density map and (top right) the CC values between these models and the opposite half maps (i.e. models built into half map 1 are checked against half map 2). CC values for each model are shown as light blue lines, the average over all top scoring models is plotted as a thick blue line. The overall CC values reported are averaged over the top ten scoring models for each half map. (Bottom right) auto-DRRAFTER modeling convergence computed between models refined into separate half maps. (A) *Tetrahymena* ribozyme, (B) hc16 product, (C) hc16, (D) *V. cholerae* glycine riboswitch with glycine, (E) *V. cholerae* glycine riboswitch, (F) *F. nucleatum* glycine riboswitch with glycine, (G) *F. nucleatum* glycine riboswitch, (H) eterna3D-JR_1, (I) spinach-TTR-3, (J) ATP-TTR-3 with AMP, (K) ATP-TTR-3, (L) SAM-IV riboswitch with SAM, and (M) SAM-IV riboswitch.

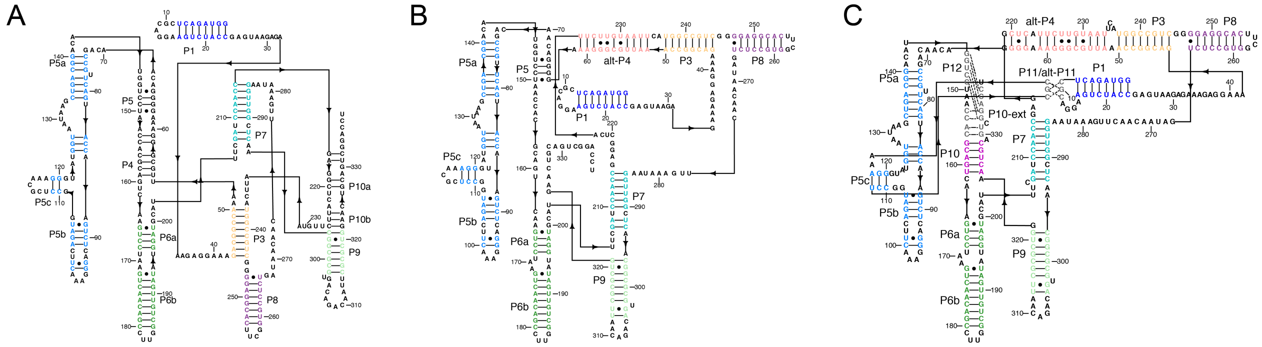


**Fig. S21.**

Determining the hc16 secondary structure. (A) The previously proposed hc16 secondary structure (*4*). (B) M2-seq and mutate-map-rescue experiments suggest that hc16 contains alt-P4 rather than P4. (C) Additional modeling and experiments suggest that alt-P4 is extended, a pseudoknot is formed between the hairpin loops of P5c and P1, P5 is not formed, and P10 is formed.

**Table S1.**

Cryo-EM data collection and processing.

| System | Molecular weight (nt/kDa) | Microscope | Voltage (kV) | Pixel size (Å) | Total Dose (e-/Å$^2$) | Phase plate | No. micrographs | No. initial particles | No. final particles | Resolution (0.143 FSC, Å) |
|---|---|---|---|---|---|---|---|---|---|---|
| *Tetrahymena* ribozyme | 388/125 | Talos Arctica | 200 | 1.07 | 30 | Yes | 752 | 233,365 | 74,621 | 6.8 |
| hc16 product | 349/112 | Talos Arctica | 200 | 1.07 | 34 | Yes | 1,414 | 287,851 | 29,191 | 10 |
| hc16 | 338/108 | Talos Arctica | 200 | 1.07 | 34 | Yes | 1,012 | 228,610 | 21,263 | 10 |
| ScaRNA6 | 265/85 | Titan Krios | 300 | 1.07 | 54 | No | 860 | 36,308 | 19,363 | N/A |
| *V. cholerae* glycine riboswitch with glycine | 231/79 | Titan Krios | 300 | 0.65 | 45 | No | 4,700 | 509,061 | 193,317 | 5.7 |
| *V. cholerae* glycine riboswitch (apo) | 231/79 | Titan Krios | 300 | 1.06 | 10.6 | No | 6,600 | 792,328 | 230,891 | 4.8 |
| RB1 5' UTR | 189/61 | Talos Arctica | 200 | 1.07 | 54 | Yes | 1,050 | 51,618 | 27,143 | N/A |
| 24-3 | 180/57 | Titan Krios | 300 | 1.06 | 45.6 | No | 2,260 | 353,447 | 123,217 | N/A |
| *F. nucleatum* glycine riboswitch with glycine | 171/55 | Talos Arctica | 200 | 1.07 | 30 | Yes | 1,311 | 677,706 | 35,578 | 7.4 |
| *F. nucleatum* glycine riboswitch (apo) | 171/55 | Talos Arctica | 200 | 1.07 | 30 | Yes | 959 | 380,061 | 20,269 | 10 |
| U1 snRNA | 168/54 | Talos Arctica | 200 | 1.07 | 54 | No | 880 | 46,983 | 18,277 | N/A |
| Eterna3D-JR_1 | 135/43 | Talos Arctica | 200 | 1.07 | 55.2 | Yes | 600 | 77,647 | 51,690 | 14 |
| Spinach-TTR-3 | 130/42 | Talos Arctica | 200 | 1.07 | 55.2 | Yes | 1,100 | 169,120 | 67,474 | 12 |
| ATP-TTR-3 with AMP | 130/42 | Talos Arctica | 200 | 1.07 | 30 | Yes | 934 | 183,317 | 39,136 | 10 |
| ATP-TTR-3 (apo) | 130/42 | Talos Arctica | 200 | 1.07 | 30 | Yes | 977 | 445,539 | 71.045 | 10 |
| SAM IV riboswitch (with SAM) | 119/40 | Titan Krios | 300 | 1.06 | 45.6 | No | 3,200 | 849,704 | 225,303 | 4.8 |
| SAM IV riboswitch (apo) | 119/40 | Titan Krios | 300 | 1.06 | 45.6 | No | 2,860 | 854,888 | 260,244 | 4.7 |
| Downstream peptide riboswitch with glutamine | 62/20 | Talos Arctica | 200 | 1.07 | 30 | Yes | 983 | 514,337 | 32,797 | N/A |

**Table S2.**

Accuracy of auto-DRRAFTER models built into simulated density maps.

| System | No. nucleotides | Convergence (Å) | Mean RMSD (Å) | Min RMSD (Å) | CC | No. modelin g rounds | Final round modeling performed ? |
|---|---|---|---|---|---|---|---|
| THF riboswitch (3SUX) | 101 | 3.03 | 4.02 | 3.81 | 0.94 | 3 | Yes |
| Bacterial SRP Alu domain (4WFL) | 105 | 2.10 | 2.48 | 2.25 | 0.97 | 4 | Yes |
| FMN riboswitch (3F2Q) | 112 | 12.12 | 9.17 | 5.92 | 0.85 | 9 | Yes |
| SAM-I riboswitch (4KQY) | 119 | 2.97 | 3.26 | 2.98 | 0.95 | 4 | Yes |
| c-di-AMP riboswitch (4QK8) | 122 | 2.93 | 3.33 | 3.00 | 0.95 | 4 | Yes |
| *Tetrahymena* ribozyme P4-P6 domain (1GID) | 158 | 6.36 | 6.34 | 5.24 | 0.91 | 7 | Yes |
| Lysine riboswitch (3DIL) | 174 | 4.21 | 4.93 | 4.51 | 0.95 | 5 | Yes |
| Lariat capping ribozyme (4P8Z) | 188 | 6.92 | 11.07 | 9.99 | 0.85 | 8 | Yes |

**Table S3.**

Convergence and CC for best-case models.

| System | No. nts | Map resolution (Å) | Convergence (Å) | Convergence half 1 | Convergence half 2 | CC full map | CC half map 1 | CC half map 2 | No. modeling rounds |
|---|---|---|---|---|---|---|---|---|---|
| *Tetrahymena* ribozyme | 388 | 6.8 | 2.4 | 2.4 | 2.4 | 0.80 | 0.83 | 0.83 | 3 |
| hc16 product | 349 | 10.0 | 6.4 | 6.3 | 6.6 | 0.79 | 0.84 | 0.84 | 3 |
| hc16 product conformation 2 | 349 | 10.0 | 5.4 | 5.4 | 5.5 | 0.79 | 0.80 | 0.80 | 3 |
| hc16 | 338 | 10.0 | 6.2 | 6.2 | 6.1 | 0.81 | 0.82 | 0.82 | 3 |
| *V. cholerae* glycine riboswitch with glycine | 231 | 5.7 | 2.4 | 2.4 | 2.4 | 0.73 | 0.74 | 0.74 | 5 |
| *V. cholerae* glycine riboswitch apo | 231 | 4.8 | 1.4 | 1.4 | 1.3 | 0.67 | 0.69 | 0.69 | 4 |
| *F. nucleatum* glycine riboswitch with glycine | 171 | 7.4 | 1.6 | 1.5 | 1.6 | 0.80 | 0.80 | 0.80 | 3 |
| *F. nucleatum* glycine riboswitch apo | 171 | 10.0 | 1.8 | 1.9 | 1.7 | 0.89 | 0.88 | 0.88 | 3 |
| Eterna3D-JR_1 | 135 | 14.0 | 2.5 | 2.5 | 2.5 | 0.83 | 0.83 | 0.84 | 10 |
| Spinach-TTR-3 | 130 | 12.0 | 2.9 | 2.9 | 3.0 | 0.87 | 0.87 | 0.87 | 6 |
| ATP-TTR-3 with AMP | 130 | 10.0 | 2.2 | 1.9 | 2.4 | 0.79 | 0.83 | 0.84 | 3 |
| ATP-TTR-3 apo | 130 | 10.0 | 2.1 | 2.2 | 1.7 | 0.62 | 0.81 | 0.81 | 3 |
| SAM-IV riboswitch with SAM | 119 | 4.8 | 1.9 | 1.9 | 1.9 | 0.76 | 0.77 | 0.78 | 3 |
| SAM-IV riboswitch apo | 119 | 4.7 | 2.1 | 2.1 | 2.0 | 0.71 | 0.71 | 0.70 | 3 |

**Table S4.**

Assessing automated models: convergence, CC, and RMSD to best-case models.

| System | No. nts | Map resolution (Å) | Convergence | Convergence half-1 (Å) | Convergence half-2 (Å) | Mean RMSD (Å) | Min. RMSD (Å) | CC full | CC half-1 | CC half-2 | No. modeling rounds | Final round modeling performed? |
|---|---|---|---|---|---|---|---|---|---|---|---|---|
| *Tetrahymena* ribozyme | 388 | 6.8 | 56.6 | N/A | N/A | 38.6 | 38.5 | 0.51 | - | - | 7 | No |
| hc16 product | 349 | 10.0 | 56.2 | N/A | N/A | 43.8 | 43.5 | 0.60 | - | - | 7 | No |
| hc16 | 338 | 10.0 | 49.5 | N/A | N/A | 48.0 | 47.6 | 0.63 | - | - | 7 | No |
| *V. cholerae* glycine riboswitch with glycine | 231 | 5.7 | 3.3 | 3.3 | 3.2 | 4.4 | 4.3 | 0.69 | 0.71 | 0.71 | 12 | Yes |
| *V. cholerae* glycine riboswitch | 231 | 4.8 | 3.0 | 3.1 | 2.8 | 4.0 | 3.9 | 0.66 | 0.67 | 0.67 | 5 | Yes |
| *F. nucleatum* glycine riboswitch with glycine | 171 | 7.4 | 3.3 | 3.2 | 3.4 | 4.3 | 4.2 | 0.76 | 0.77 | 0.78 | 4 | Yes |
| *F. nucleatum* glycine riboswitch apo | 171 | 10.0 | 4.0 | 4.3 | 3.6 | 8.3 | 8.2 | 0.85 | 0.86 | 0.86 | 4 | Yes |
| Eterna3D-JR_1 | 135 | 14.0 | 2.5 | 2.5 | 2.5 | - | - | 0.83 | 0.83 | 0.84 | 10 | Yes |
| Spinach-TTR-3 | 130 | 12.0 | 3.5 | 3.5 | 3.6 | 7.9 | 7.7 | 0.88 | 0.88 | 0.88 | 9 | Yes |
| ATP-TTR-3 with AMP | 130 | 10.0 | 2.0 | 1.9 | 2.0 | 3.8 | 3.5 | 0.8 | 0.84 | 0.85 | 3 | Yes |
| ATP-TTR-3 apo | 130 | 10.0 | 2.0 | 1.9 | 1.9 | 4.2 | 4.0 | 0.63 | 0.82 | 0.81 | 4 | Yes |
| SAM-IV riboswitch with SAM | 119 | 4.8 | 25.4 | N/A | N/A | 21.5 | 21.4 | 0.6 | - | - | 7 | No |
| SAM-IV riboswitch apo | 119 | 4.7 | 3.6 | 3.5 | 3.5 | 5.6 | 5.6 | 0.66 | 0.67 | 0.66 | 5 | Yes |

**Table S5.**

Nucleotides with convergence above 10 Å or CC below 0.5.

| System | Nucleotides with convergence > 10 Å | Nucleotides with CC < 0.5 |
|---|---|---|
| *Tetrahymena* ribozyme best-case | 22 | none |
| *Tetrahymena* ribozyme automated | 22-409 | 22, 23, 24, 25, 30, 31, 32, 33, 34, 35, 36, 37, 38, 39, 40, 41, 42, 43, 44, 45, 46, 47, 48, 49, 50, 51, 52, 53, 54, 55, 56, 57, 102, 103, 104, 105, 110, 111, 112, 113, 114, 115, 116, 149, 150, 151, 152, 153, 154, 169, 170, 171, 172, 173, 175, 176, 177, 178, 184, 207, 208, 209, 210, 211, 212, 213, 214, 215, 216, 217, 218, 252, 253, 254, 255, 256, 257, 258, 259, 260, 261, 262, 263, 264, 288, 289, 290, 305, 306, 307, 308, 313, 314, 315, 316, 317, 318, 319, 320, 321, 322, 323, 324, 325, 326, 327, 328, 329, 330, 331, 332, 333, 334, 335, 336, 337, 338, 349, 350, 351, 360, 362, 363, 364, 365, 366, 367, 368, 369, 370, 371, 381, 382, 383, 384, 385, 386, 387, 388, 391, 392, 393, 394, 395, 396, 397 |
| hc16 product best-case | 12, 13, 14, 69, 70, 71, 72, 129, 130, 131, 147, 148, 149, 150, 151, 152, 153, 154, 203, 269, 270, 271, 272, 273, 274, 275, 276, 277, 278, 279, 332, 333, 334, 335, 336, 337, 338 | none |
| hc16 product automated | -10-338 | 170, 171, 181, 182, 183, 184, 271, 272, 274 |
| hc16 product conformation 2 best-case | 12, 13, 36, 37, 38, 39, 40, 70, 71, 116, 129, 130, 131, 203, 204, 269, 270, 271, 272, 273, 274, 275, 276, 277, 278, 279, 280 | none |
| hc16 best-case | 12, 13, 29, 30, 31, 32, 33, 34, 35, 36, 37, 38, 68, 69, 70, 71, 72, 128, 129, 130, 217, 268, 269, 270, 271, 272, 273, 274, 275, 276, 277, 278, 279, 280, 281, 282, 336, 337, 338 | none |
| hc16 automated | 1-338 | 46, 47, 241, 242, 243, 244, 245, 246, 247, 248, 264, 265, 266, 271, 278, 279, 280, 281, 282, 283, 284, 285, 286, 338 |
| *V. cholerae* glycine riboswitch with glycine best-case | none | 104 |

| | | |
|---|---|---|
| *V. cholerae* glycine riboswitch with glycine automated | -4 | none |
| *V. cholerae* glycine riboswitch apo best-case | none | 103, 104 |
| *V. cholerae* glycine riboswitch apo automated | none | 101, 102, 103, 104, 105 |
| *F. nucleatum* glycine riboswitch with glycine best-case | none | none |
| *F. nucleatum* glycine riboswitch with glycine automated | none | none |
| *F. nucleatum* glycine riboswitch apo best-case | none | none |
| *F. nucleatum* glycine riboswitch apo automated | none | none |
| Eterna3D-JR_1 automated | none | none |
| Spinach-TTR-3 best-case | none | none |
| Spinach-TTR-3 automated | 110, 111 | none |
| ATP-TTR-3 with AMP best-case | none | none |
| ATP-TTR-3 with AMP automated | none | none |
| ATP-TTR-3 apo best-case | none | none |
| ATP-TTR-3 apo automated | none | none |
| SAM-IV riboswitch with SAM best-case | none | none |
| SAM-IV riboswitch with SAM automated | 1, 2, 3, 4, 5, 6, 7, 8, 9, 10, 11, 12, 13, 14, 15, 16, 17, 18, 19, 20, 21, 22, 23, 24, 25, 26, 27, 28, 29, 30, 31, 32, 33, 34, 35, 36, 38, 40, 41, 42, 43, 44, 45, 46, 47, 48, 49, 50, 51, 52, 53, 54, 55, 56, 57, 58, 59, 60, 61, 62, 63, 64, 65, 66, 67, 68, 69, 70, 71, 72, 73, 74, 75, 76, 77, 78, 79, 80, 81, 82, 83, 84, 85, 86, 87, 88, 89, 90, 91, 92, 93, 94, 95, 96, 97, 98, 99, 100, 101, 102, 103, 104, 105, 106, 107, 108, 109, 110, 111, 112, 113, 114, 115, 116, 117, 118, 119 | 50, 51, 52, 53, 54, 55, 56, 96, 97 |
| SAM-IV riboswitch apo best-case | none | none |
| SAM-IV riboswitch apo automated | 113, 114, 115 | none |

**Table S6. (separate file)**
DNA and RNA sequences.
